# Mycothiazole enables structural characterization of entire ubiquinone-accessing tunnel in mitochondrial respiratory complex I

**DOI:** 10.64898/2026.10.01.756109

**Authors:** Shinsuke Akisada, Ryohei Otani, Takahiro Masuya, Shuji Miyako, Hiroki Saito, Kaoru Mitsuoka, Hideto Miyoshi, Ken Yokoyama, Masatoshi Murai

## Abstract

Mitochondrial NADH-ubiquinone (UQ) oxidoreductase (respiratory complex I) reduces UQ within its long tunnel-like cavity, which is coupled to triggering proton translocation at the membrane domain. Although mammalian complex I adopts distinct “open” and “closed” conformations, the architecture of the UQ-accessing tunnel in the open conformation remains poorly defined owing to partial disorder of the UQ catalytic region near Fe-S cluster N2 in the tunnel. Here, we applied cryo-electron microscopy (EM) single particle analysis to determine the structures of complex I in bovine heart submitochondrial particles in the presence or absence of a mycothiazole-type inhibitor. Preceding photoaffinity labeling experiments using a synthetic photoreactive derivative ([^125^I]pMT) revealed that mycothiazole specifically labels the 49-kDa and ND1 subunits, both of which form a part of the UQ-accessing tunnel. Cryo-EM analysis demonstrated that bound mycothiazole reorganizes and stabilizes the UQ catalytic region in both open and closed conformations, enabling the first complete structural modeling of the entire UQ-accessing tunnel in the open conformation. Unexpectedly, in the presence of bound mycothiazole, loops of the PSST subunit forming the catalytic region in the tunnel adopted closed conformation-like features, whereas open conformation-like features were determined for the transmembrane helices of ND1 and ND6 subunits. The present study revealed that bound mycothiazole can reorganize the UQ catalytic region in the UQ-accessing tunnel and lead to a hybrid conformational state beyond the canonical open–*versus*–closed conformational model.

**Significance Statement:** Respiratory complex I is a central energy-converting enzyme that couples ubiquinone (UQ) reduction to proton translocation across the inner mitochondrial membrane. Mammalian complex I adopts distinct open and closed conformations, but the functional significance of the open conformation remains unresolved. Structural characterization of the UQ-reduction region in the open conformation has been limited by disorder of key elements surrounding the catalytic site. Using mycothiazole as a molecular probe, we stabilized and visualized the entire UQ-accessing tunnel in the open conformation. Unexpectedly, mycothiazole reorganized the catalytic region in both conformations into a common hybrid architecture, revealing that structural elements around the UQ catalytic site and those linked to proton translocation can rearrange independently.

## Introduction

Proton-translocating NADH-ubiquinone (UQ) oxidoreductase, known as respiratory complex I, catalyzes the initial step of the mitochondrial respiratory chain. It couples electron transfer from NADH to UQ with proton translocation across the inner mitochondrial membrane, thereby generating the proton-motive force required for ATP synthesis and substrate transport [*1–4*]. Complex I is composed of two distinct domains: a hydrophilic domain, which extends into the mitochondrial matrix and mediates electron transfer, and a membrane domain embedded within the inner mitochondrial membrane, which is responsible for proton translocation. As complex I plays a central role in cellular energy metabolism, dysfunction of this enzyme leads to severe neurodegenerative and metabolic disorders, and is also implicated in ischemia-reperfusion injury, a major form of oxidative tissue damage [*5,6*].

X-ray crystallographic and cryo-electron microscopy (cryo-EM) structures of complex I from various organisms have provided important insights into the redox-driven proton-pumping mechanism of the enzyme [*7–13*]. The outcomes of these studies have led to the consensus that structural and electrostatic rearrangements induced by UQ reduction at the interfacial region between the hydrophilic and membrane domains transmit to the membrane subunits to trigger proton translocation [*1–4,7*]. The UQ-reaction site is highly conserved among organisms and has been identified as an unusually long and narrow tunnel-like cavity, termed the UQ-accessing tunnel (approximately 30 Å long and 5 Å in diameter at the entrance). It is generally considered that UQ molecules enter and transit the tunnel to be reduced at its “top” near a terminal iron-sulfur cluster N2 [*1–4*].

The unusual architecture of the UQ-accessing tunnel represents one of the most enigmatic structural features of complex I. Longstanding debate has focused on how UQ transits close to the Fe-S cluster N2 and moves back from the tunnel after the UQ head-ring is reduced, and how this UQ reduction process triggers proton translocation in the membrane domain. Several mechanistic models have been proposed based on high-resolution structures of complex I in combination with biochemical [*14–16*] and computational [*17–22*] studies. However, the molecular details underlying the energy-coupling mechanism remain incompletely understood.

Recent cryo-EM studies have also disclosed the binding positions of a variety of inhibitors in mammalian complex I [*12, 23–28*]. These structural studies not only show that the inhibitors interfere with the UQ reduction in a competitive manner against a substrate UQ, but also that various inhibitors compete with each other to occupy the tunnel’s interior. Nevertheless, our previous biochemical studies are difficult to reconcile with these scenarios. For example, photoaffinity labeling experiments using photoreactive amiloride-type inhibitors showed that some amilorides specifically label the C-terminal region (Thr227−Lys283) of the 39-kDa subunit, which does not comprise part of the canonical UQ-accessing tunnel [*29*]. However, this labeling is completely suppressed by excess rotenone and piericidin A, which bind to the interior of the tunnel [*12,23*]. Furthermore, photoaffinity labeling experiments using S1QELs (small molecules that suppress superoxide production from complex I during reverse electron transport [*30*]) and IACS-010759 (complex I inhibitor originally developed as suppressor of hypoxia-inducible factor-1α [*31,32*]) showed that these inhibitors bind to the ND1 subunit, but their binding was not suppressed by rotenone or piericidin A. Based on these results along with other findings [*33,34*], we proposed that the *matrix-side* interface region of flexible loops of the 49-kDa (β1–β2), ND1 (TMHs 5–6), and ND3 (TMHs 1–2) subunits may be a candidate for the alternative route(s) of ligand molecules to the catalytic site at the “top” of the tunnel. Cryo-EM studies on short-chain UQ_1_-bound porcine complex I [*26*] and medicinal biguanides-bound bovine complex I [*23,28*], and an MD simulation study on UQ_1_ reaction in the tunnel [*35*] also suggested marked flexibility of this matrix-side interface region of the tunnel.

Mycothiazole (Figure 1) was discovered as a suppressor of hypoxia-inducible factor-1α via targeting mitochondrial complex I [*36*]. Here, we investigated the action manner of mycothiazole in complex I using a photoaffinity labeling technique and cryo-EM analyses with bovine heart submitochondrial particles (SMPs). The photoaffinity labeling experiments identified the 49-kDa and ND1 subunits, two components forming the UQ-accessing tunnel, as the major labeled regions. Importantly, tunnel-forming elements hitherto unresolved in the open conformation became ordered in the presence of bound mycothiazole, enabling complete modeling of the tunnel’s architecture. In addition, these elements adopted neither the canonical open nor closed arrangement but instead formed a hybrid structural state in which closed- and open-conformation-like features coexist around the catalytic site in the tunnel. These findings provide the first complete structural characterization of the UQ-accessing tunnel in the open conformation, and reveal local conformational plasticity beyond the canonical open–closed framework.

**Figure 1.**
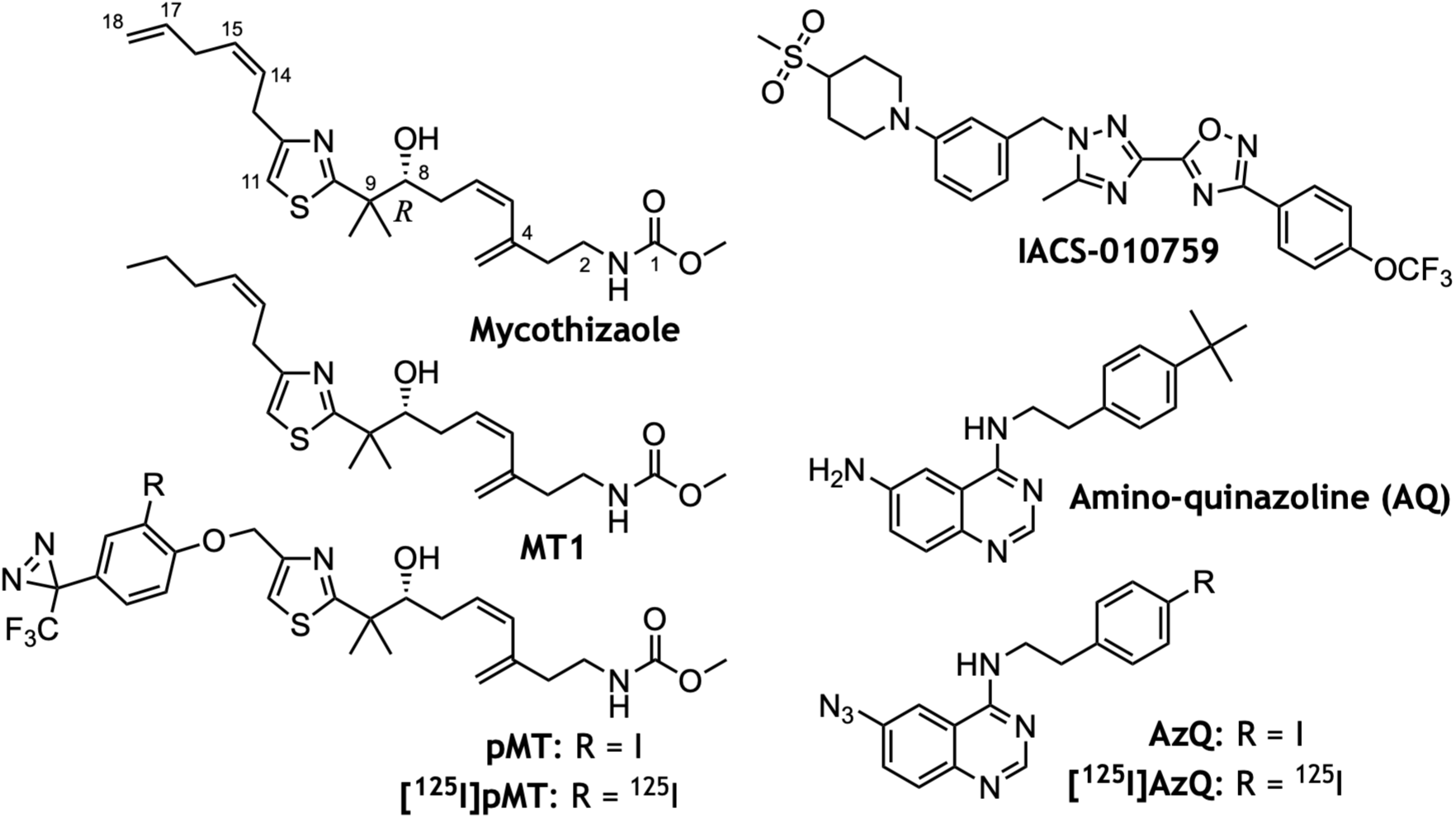
Structures of complex I inhibitors used or discussed in this study. Natural mycothiazole was unavailable; therefore, the synthetic analogue MT1 was synthesized and used throughout this study.

## Results

### Synthetic mycothiazole inhibits bovine complex I at ubiquinone-binding site

A characteristic structural feature of mycothiazole is a central thiazole ring flanked by two acyclic polyketide chains: one is an alkyl chain containing two double bonds at C14/C15 and C17/C18, and the other is an alkyl methylcarbamate moiety containing a conjugated diene system and an (*R*)-configured hydroxyl group at C8 (Figure 1). To identify the binding site of mycothiazole in complex I, we sought to develop its photoreactive derivative. Based on structure–activity relationships reported for complex I inhibitors possessing similar structural frameworks [*37,38*], we considered that chemical modifications of the alkyl side chain attached to the thiazole ring would have minimal effects on the inhibitory activity. Accordingly, we designed a synthetic route to introduce a photoreactive group into this moiety while preserving the remaining structural frameworks of mycothiazole (Figure 1, Schemes S1–S3). Because of difficulty synthesizing the two double bonds at C14/C15 and C17/C18 of natural mycothiazol, we prepared MT1 (Figure 1) and used it as a reference compound of mycothiazole throughout this study.

Morgan et al. [*36*] identified complex I as the molecular target of mycothiazole by monitoring oxygen consumption in digitonin-permeabilized T47D cells. To verify this, we directly investigated the effect of MT1 on complex I activity using bovine heart submitochondrial particles (SMPs). MT1 inhibited the NADH-UQ_1_ oxidoreductase activity with an IC_50_ value of 27 (± 5) nM (Figure 2A). The result confirms that mycothiazole is an inhibitor of complex I, although its inhibitory potency is approximately one order of magnitude weaker than that of traditional complex I inhibitors, such as rotenoids, piericidins, and acetogenins, which exhibit inhibition at single-digit nanomolar IC_50_ levels (1–3 nM) under the same experimental conditions [*39,40*].

**Figure 2.**
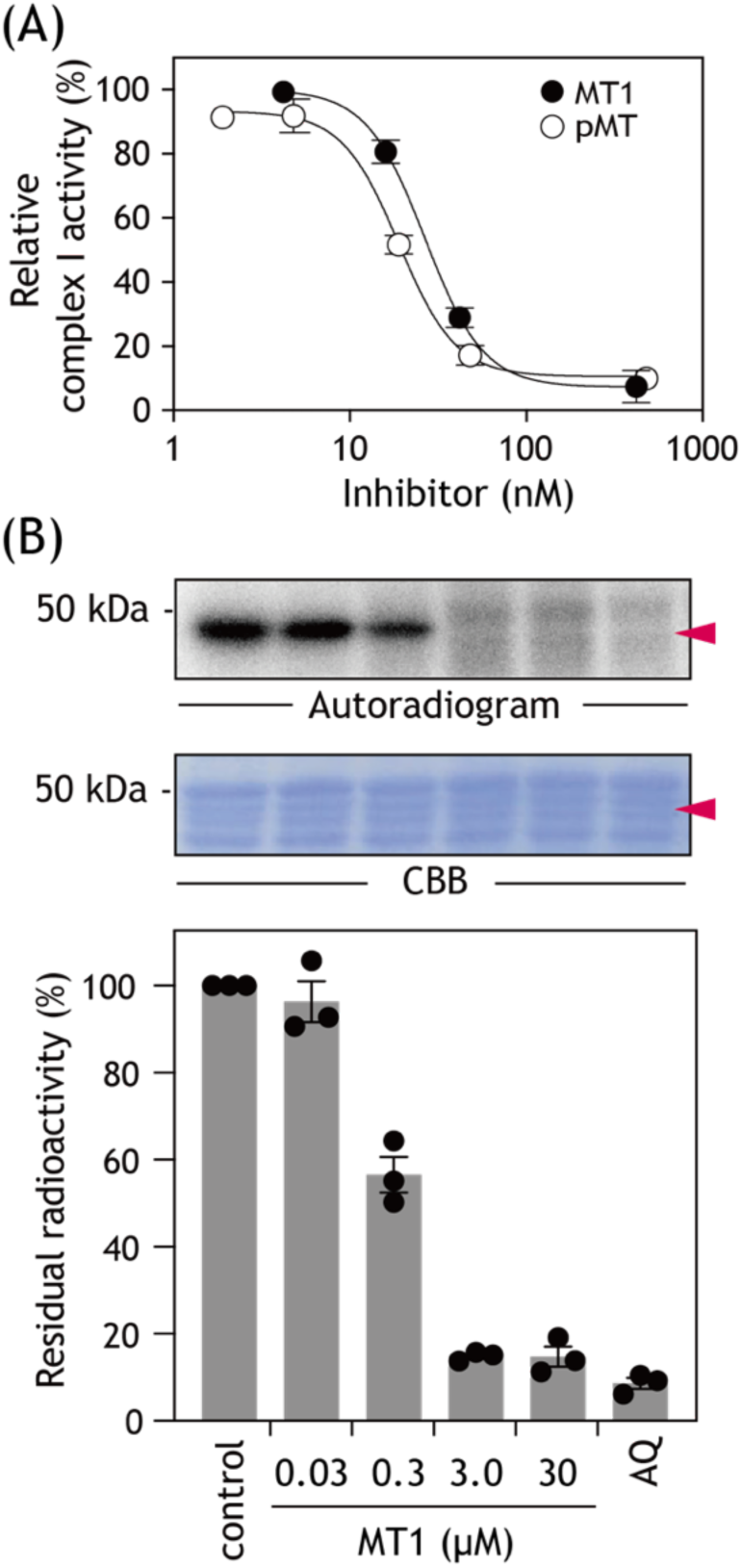
Synthetic mycothiazole inhibits mammalian complex I and competes for ubiquinone-binding site. (A) Effects of MT1 and pMT on NADH-UQ_1_ oxidoreductase activity. Bovine SMPs (30 µg protein/mL) were incubated with increasing concentrations of MT1 (*closed circles*) or pMT (*open circles*), and NADH-UQ_1_ oxidoreductase activity was measured. Data are presented as the mean ± SD (*n* = 3). (B) MT1 suppresses labeling of 49-kDa subunit by the photoreactive quinazoline [¹²⁵I]AzQ. Bovine SMPs (2.0 mg protein/mL) were UV-irradiated with 5.0 nM [¹²⁵I]AzQ in the presence of increasing concentrations of MT1. Samples were separated by SDS-PAGE, followed by CBB staining and autoradiography. *Upper panel,* autoradiogram of the 49-kDa subunit labeled by [¹²⁵I]AzQ in the absence (control) or presence of the indicated concentrations of MT1. *Middle panel,* CBB staining of the corresponding region showing equal protein loading. *Lower panel,* quantification of residual radioactivity associated with the 49-kDa subunit. Radioactivity was normalized to the control sample (100%). Approximately 40 µg of protein was loaded per lane. Data are presented as the mean ± SE (*n* = 3). The rightmost lane shows SMPs UV-irradiated in the presence of amino-quinazoline (AQ), the non-photoreactive parent compound of AzQ, as a reference competitor.

An X-ray crystallographic study on *Yallowia lipolytica* complex I showed that quinazoline-type inhibitors bind to the “top” of the UQ-accessing tunnel [*8*]. Next, we examined whether MT1 suppresses the binding of a photoreactive quinazoline [^125^I]AzQ (Figure 1) to the UQ-accessing tunnel of bovine complex I (Figure 2B). Previous photoaffinity-labeling studies showed that [^125^I]AzQ labels the interface between the N-terminal region of the 49-kDa subunit (Asp41–Arg63) and third matrix-side loop connecting transmembrane helices (TMHs) 5 and 6 of ND1 (Asp199–Lys262), both of which contribute to forming the UQ-accessing tunnel (Figure S1) [*41,42*]. Although both regions are labeled, most of the incorporated radioactivity is associated with the 49-kDa subunit [*41,42*]. When SMPs were labeled with 5.0 nM [^125^I]AzQ in the presence of different concentrations of MT1, labeling of the 49-kDa subunit was suppressed in a concentration-dependent manner, with nearly complete suppression observed at 3.0 µM MT1 (Figure 2B). The results suggest that MT1 occupies the UQ-accessing tunnel in a manner similar to that of classical complex I inhibitors.

### Photoaffinity labeling identifies 49-kDa and ND1 subunits as binding sites of mycothiazole

To assess the binding site of mycothiazole in bovine complex I, we synthesized [^125^I]pMT (Figure 1), a photoreactive mycothiazole derivative containing a photoreactive diazirine and radioisotope (^125^I) at the terminal end of the molecule. The IC_50_ value of the corresponding non-radioactive derivative (pMT) was 19 nM (Figure 2A), being comparable to that of the parent compound MT1 above. Bovine SMPs were photoaffinity-labeled with 5.0 nM [^125^I]pMT, separated by BN-PAGE, and the complex I band was subsequently subjected to doubled SDS-PAGE to resolve the major subunits [*43–45*]. Although the incorporated radioactivity was relatively weak compared with that observed in previous photoaffinity-labeling studies [*41,46,47*], likely reflecting the lower binding affinity of [^125^I]pMT in terms of the IC_50_ value, we reproducibly detected radioactive spots corresponding to the 49-kDa and ND1 subunits (Figure 3A), both of which contribute to forming the UQ-accessing tunnel. Although the ND5 subunit located in the membrane domain was also labeled, we will discuss the specificity of this labeling below.

**Figure 3.**
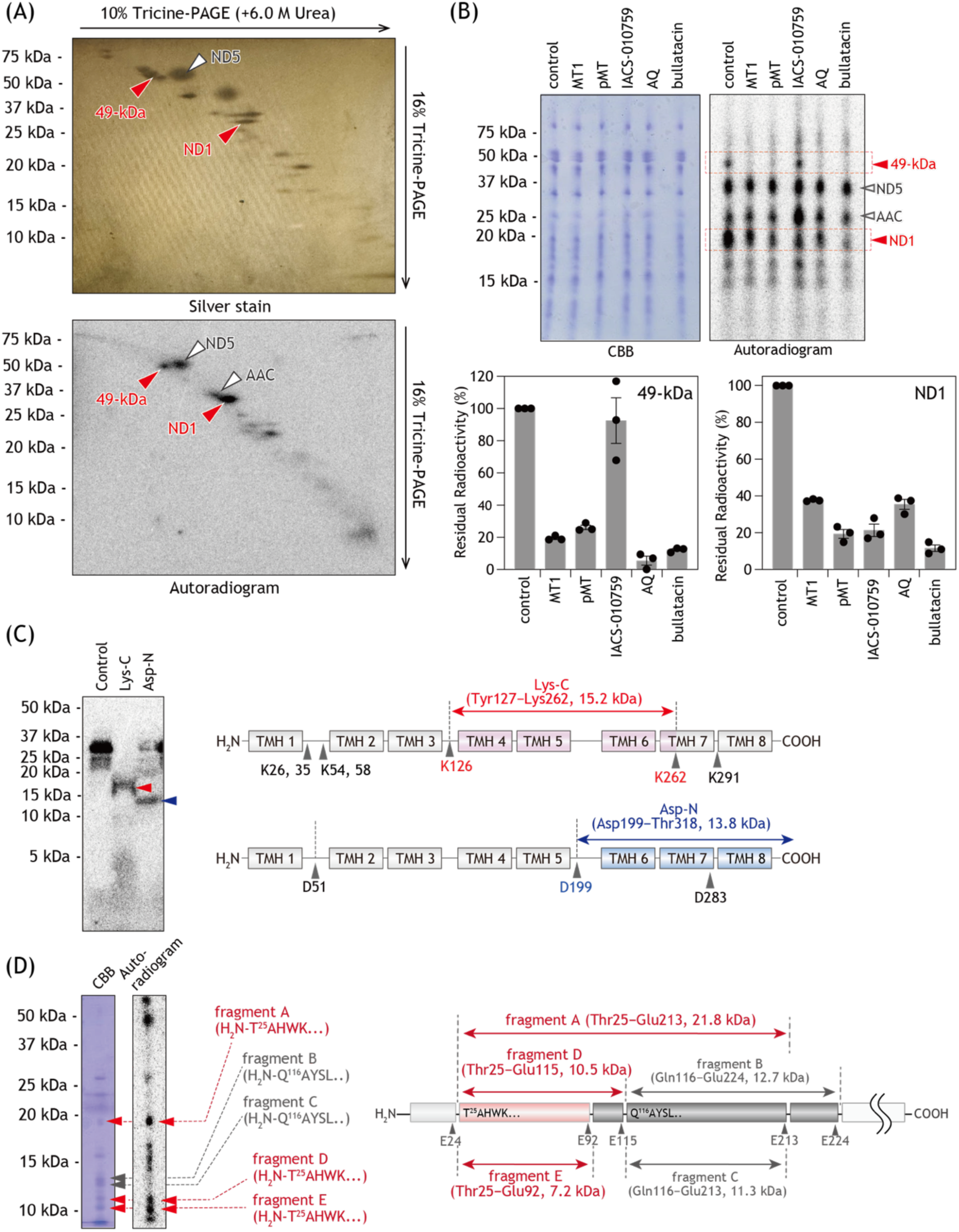
Localization of labeled regions in 49-kDa and ND1 subunits of bovine complex. **I.** (A) Identification of the subunits labeled by [¹²⁵I]pMT. Bovine SMPs (2.0 mg/mL, 100 µL) were labeled with [¹²⁵I]pMT (5.0 nM), and complex I was isolated by BN-PAGE. The isolated complex I was subsequently separated by dSDS-PAGE, followed by silver staining (*upper*) and autoradiography (*lower*). The labeled 49-kDa and ND1 subunits are indicated by red arrowheads. (B) Competition of labeling of 49-kDa and ND1 subunits by [¹²⁵I]pMT. SMPs (2.0 mg/mL) were labeled with [¹²⁵I]pMT (5.0 nM) in the presence of an excess of the indicated complex I inhibitors (5.0 µM, 1,000 molar-fold). The samples were separated by BN-PAGE, and the complex I band was excised and subsequently separated by 10% Tricine gel (10% T, 3% C) containing 6.0 M urea. The gel was stained with CBB (*left*), and radioactivity associated with the 49-kDa and ND1 subunits was detected and quantified by autoradiography (*right*). Lower panels show quantification of residual radioactivity associated with the 49-kDa and ND1 subunits in the presence of the indicated competitors. Radioactivity was normalized to the control sample (100%). Data are presented as the mean ± SE (*n* = 3). We note that [^125^I]pMT also labeled the ADP/ATP carrier (indicated as *“AAC”* in *panels A and B*), as previously observed in our photoaffinity labeling studies using photoreactive complex I inhibitors [*29,30,47*]. (C) Localization of the labeled region in the ND1 subunit. The ND1 band isolated from the gel shown in (B) was digested with Lys-C or Asp-N, and the digests were separated by 16% Tricine SDS-PAGE (16% T, 6% C). Autoradiograms of the resulting fragments (*left*) and corresponding proteolytic maps based on the theoretical cleavage sites (*right*) are shown. (D) Localization of the labeled region in the 49-kDa subunit. The 49-kDa band isolated from the gel presented in (B) was subjected to Cleveland mapping. CBB staining and autoradiography of the resulting fragments are shown on the *left*. The corresponding proteolytic map, constructed from fragment assignments previously determined by Edman degradation [*45*], is shown on the *right*.

When SMPs were labeled with 5.0 nM [^125^I]pMT in the presence of mycothiazole derivatives (MT1 and pMT) or other traditional complex I inhibitors (IACS-010759, amino-quinazoline (AQ), and bullatacin), labeling of the 49-kDa and ND1 subunits was suppressed to approximately 20 and 35% of the control level, respectively. However, labeling of ND5 was not suppressed by any of the inhibitors tested (Figure 3B). Note that the significance of ND5 labeling remains unclear, as discussed later. Taken together, the results support the suggestion that the 49-kDa and ND1 subunits, both of which contribute to formation of the UQ-access tunnel [*1–4*], are the specific binding sites of mycothiazole. Labeling of the ND1 subunit by [^125^I]pMT was suppressed by IACS-01075 (Figure 1), whereas labeling of the 49-kDa subunit was not suppressed (Figure 3B). This different competition behavior is consistent with our previous photoaffinity-labeling study, indicating that IACS-01075 binds to the lower part of TMHs of ND1 [*32*].

We also noted that [¹²⁵I]pMT labeled a ADP/ATP carrier (AAC; Figures 3A and 3B). Similar concomitant labeling of AAC was sometimes observed in our previous photoaffinity studies using various types of complex I inhibitors possessing different photoreactive groups [*29,41,47*]. Although the precise basis for this recurrent labeling remains unclear, high abundance of AAC in the inner mitochondrial membrane may be attributable to this matter.

### Localization of [^125^I]pMT-labeled sites in ND1 and 49-kDa subunits

To identify regions labeled by [^125^I]pMT in ND1 and 49-kDa subunits, the labeled subunits were isolated from SDS gels and subjected to proteolytic digestion. Proteolytic digestion of the labeled ND1 subunit with lysyl endopeptidase (Lys-C) and endoprotease Asp-N (Asp-N) yielded radioactive bands migrating at positions corresponding to approximately 15 and 13 kDa, respectively (Figure 3C). Based on theoretical cleavage sites, these bands were assigned to the peptides Tyr127–Lys262 (15.2 kDa) and Asp199–Thr318 (13.8 kDa), respectively. These results suggest that the [^125^I]pMT-labeling site is located within the region Asp199–Lys262, which covers the large matrix-side loop connecting TMHs 5 and 6 (Figures 3C and S2). The 49 kDa subunit was digested in-gel with V8 protease, followed by peptide mapping on a 20% Tris-EDTA gel (Cleveland mapping). This digestion yielded multiple peptide fragments, including a radioactive fragment of approximately 20 kDa and two smaller radioactive fragments around 10 kDa (Figure 3D). Based on previously determined N-terminal sequences [*45*], the larger fragment was assigned to the peptide Thr25–Glu213 (21.8 kDa), whereas the two smaller fragments were assigned to its further proteolytic fragments, Thr25–Glu115 and Thr25–Glu92 (10.5 and 7.2 kDa, respectively) (Figure 3D). This region includes the loop connecting the β1 and β2 strands, which lies adjacent to the ND1 loop connecting TMHs 5 and 6 described above (Figure S2). Further proteolytic analysis of the 49-kDa subunit was unsuccessful because reproducible digestion patterns could not be obtained, possibly due to the low level of incorporated radioactivity.

### Native cryo-EM structure of mycothiazole-bound complex I

We previously determined the structures of respiratory supercomplexes in their native membrane environment through direct cryo-EM single-particle analysis of bovine heart SMPs [*48*]. This approach enabled structural determination of native mycothiazole-bound complex I. SMPs were suspended at a protein concentration of 7.0 mg/mL in buffer containing 250 mM sucrose and 10 mM Tris/HCl (pH 7.4) supplemented with 500 µM MT1. The sample was applied to a quantifoil UltraAufoil grid and flash-frozen in liquid ethane and cryo-EM micrographs were collected using Titan Krios equipped with a K3 detector. Focused classification of complex I within the I_1_III_2_IV_3_ supercomplex identified two distinct global conformations, corresponding to the canonical “closed” and “open” conformations observed in cryo-EM studies of mammalian complex I [*11,12*]. These conformations can be distinguished by the apparent angle between the hydrophilic and membrane domains and characteristic arrangements of structural elements surrounding the UQ-accessing tunnel. Hereafter, we use the terms “closed” and “open” solely to refer to these typical cryo-EM-defined conformations (Figure S3 and Table S1).

The open conformation exhibited a slightly larger angle between hydrophilic and membrane domains. The overall structures of closed and open conformations closely resembled our previously determined models without inhibitors (PDB IDs 9W2V and 9W2U, respectively, [*48*]), with RMSDs of approximately 0.57 and 0.54 Å, respectively, relative to these structures (Figure 4A).

**Figure 4.**
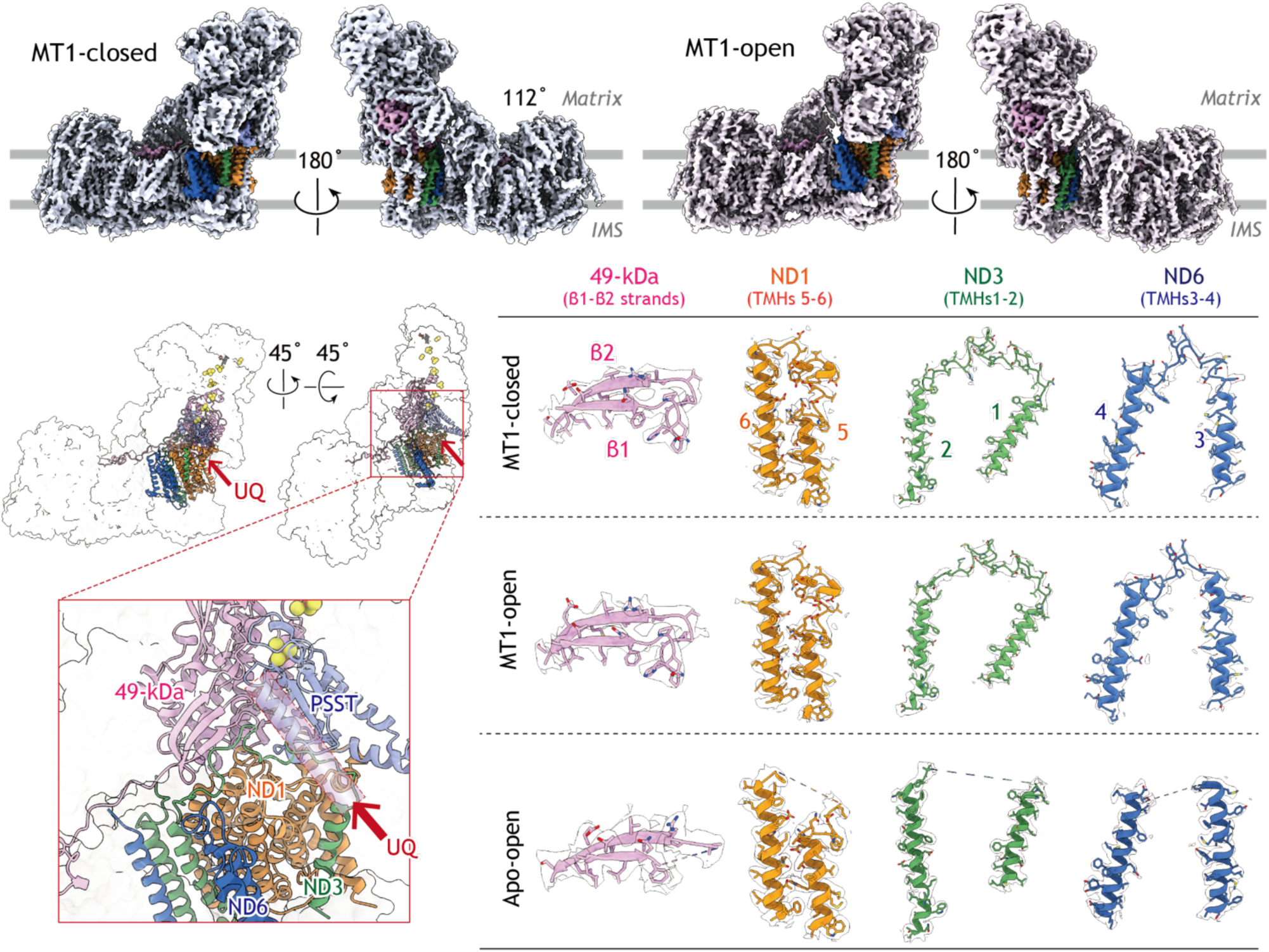
MT1 enables structural modeling of UQ-accessing tunnel in open conformation of bovine complex. **I.** (A) Cryo-EM density maps of the MT1-bound closed (*left*) and open (*right*) conformations of bovine complex I viewed parallel to the membrane plane. Two views with 180° rotation are shown for each conformation. The 49-kDa subunit (*magenta*), ND1 (*orange*), ND3 (*green*), and ND6 (*blue*) are highlighted. (B) Overall structure of the MT1-bound open conformation viewed from the membrane plane (*left*) and after successive 45° rotations (*right*). The boxed region indicates the UQ-binding site, which is enlarged below. The UQ-accessing tunnel is formed by the 49-kDa and PSST subunits together with the ND1, ND3, and ND6 subunits. The putative tunnel is indicated by the dotted rectangle. (C) Comparison of the modeled structures surrounding the UQ-accessing tunnel in the MT1-closed (*top*), MT1-open (*middle*), and apo-open (*bottom*) conformations. Cryo-EM densities are shown together with the corresponding atomic models for the β1–β2 strands of the 49-kDa subunit, ND1 TMHs5–6, ND3 TMHs1–2, and ND6 TMHs3–4. Compared with the apo-open conformation, MT1 binding enables complete modeling of these structural elements in the open conformation.

A hallmark of the open conformation of mammalian complex I is the increased flexibility of several structural elements surrounding the UQ-accessing tunnel, including four loop regions (Figures 4B and 4C): the loop connecting the β1–β2 strands of the 49 kDa subunit, that between transmembrane TMHs 5 and 6 of ND1, that between TMHs 1 and 2 of ND3, and that between TMHs 4 and 5 of ND6. In contrast, these regions are well resolved in the closed conformation, with this feature being consistently observed in structures reported by multiple research groups [*11,12*].

Interestingly, in the presence of MT1, all four loop regions were well resolved in both open and closed conformations (Figure 4C). The four loops adopted nearly identical local arrangements in the two conformations (Figure S4). Notably, in the open conformation, these loops adopted positions closely resembling those observed in the closed conformation (Figure 4C, *middle*). Together, these findings indicate that MT1 stabilizes the local structures of the UQ-accessing tunnel irrespective of the global conformation of the enzyme. Notably, the MT1-stabilized regions in the 49-kDa and ND1 subunits overlap with regions identified by the photoaffinity labeling experiments using [^125^I]pMT (Figure 3).

### Mycothiazole binding induces hybrid structural state around UQ-accessing tunnel

As MT1 stabilized structural elements surrounding the UQ-accessing tunnel in both open and closed conformations, we next examined several hallmark structural segments that distinguish the two conformations. The PSST subunit contains a pair of loops that undergo characteristic conformational rearrangements, which have been described as “*raise*” and “*flip*” transitions (Figure 5) [*12*]. In the closed conformation, the loop spanning residues Met48–Gly51 adopts the raised conformation, protruding into the UQ-access tunnel, whereas in the open conformation this loop becomes incorporated into a β-sheet. In addition, another loop comprising residues Val75–Pro80 undergoes flip transition in the open conformation, resulting in an approximately 180° rotation of the PSST-Arg77 side chain. Notably, MT1-bound complex I adopted the raised conformation in both open and closed conformations (Figure 5B), indicating that mycothiazole binding induces local rearrangement of PSST loops toward the closed conformation regardless of the global conformation of the enzyme. This pattern of PSST-loop rearrangement resembles that observed in the structure of rotenone-bound complex I [*12*].

**Figure 5.**
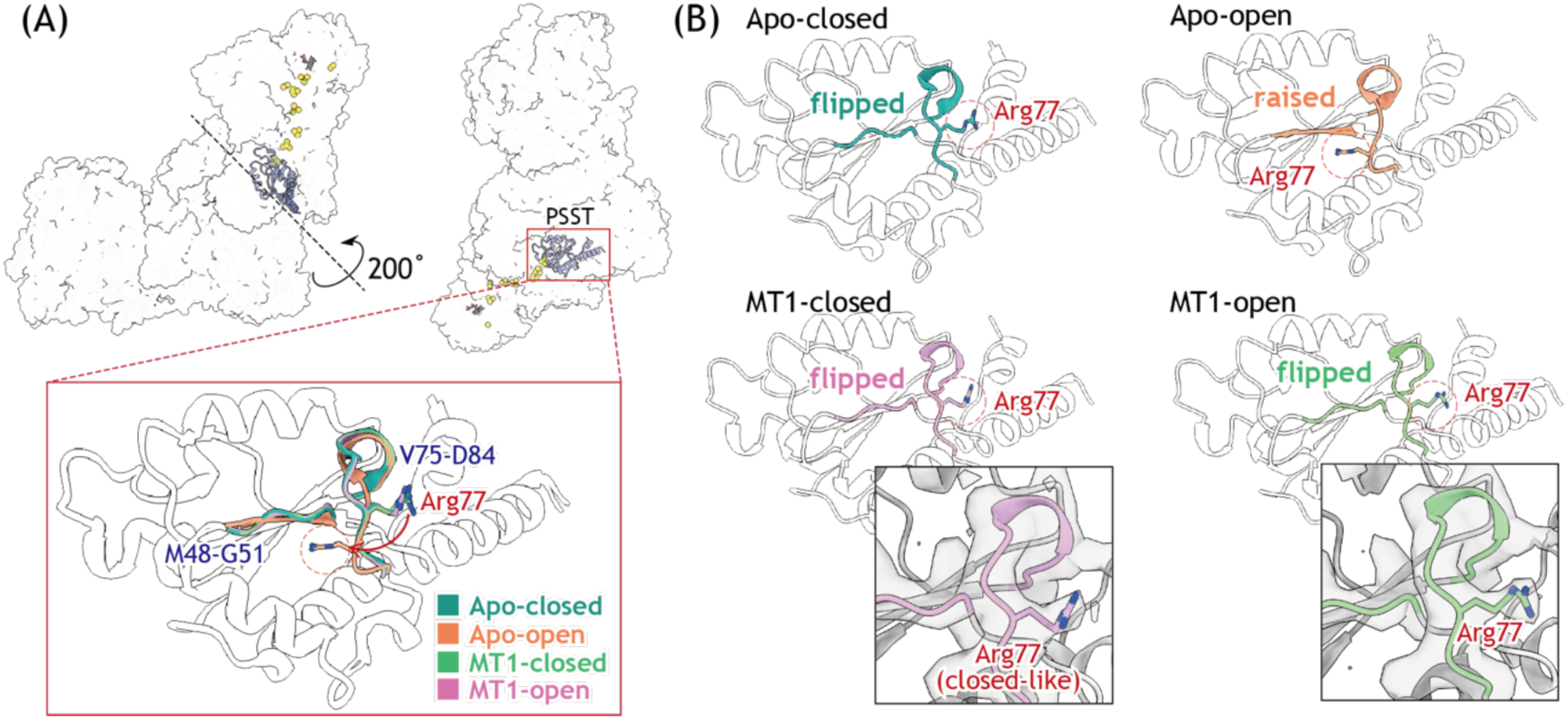
MT1 stabilizes closed-like arrangement of PSST loop in both closed and open conformations of bovine complex I. **(A)** Overall location of the PSST subunit within bovine complex I. The boxed region is enlarged to show superposition of the PSST loop in the apo-closed (*cyan*), apo-open (*orang*e), MT1-closed (*green*), and MT1-open (*magenta*) conformations. Residues Met48–Gly51, Val75–Asp84, and Arg77 are indicated. (B) Comparison of the PSST loop conformations in the apo-closed, apo-open, MT1-closed, and MT1-open structures. The apo-open structure adopts a raised conformation, whereas the apo-closed structure adopts a flipped conformation. Both MT1-bound structures exhibit the flipped (closed-like) conformation irrespective of the global conformation. Enlarged views show the cryo-EM density surrounding Arg77, supporting the modeled flipped (closed-like) conformation of Arg77.

To determine whether this local closed-state rearrangement extends to other structural segments surrounding the UQ-accessing tunnel, we next examined the conformations of ND1-TMH4 and ND6-TMH3. The UQ-accessing tunnel is connected to the proton-translocating machinery in the membrane domain through a series of protonatable residues known as the E-channel [*11,12*]. Together with conserved protonatable residues in the antiporter-like subunits ND2, ND4, and ND5, the E-channel forms a continuous water-mediated network extending along the central hydrophilic axis of the membrane domain from the UQ reaction site toward the distal region of ND5. Structural and computational studies [*12,13,22*] have proposed that this network serves as a pathway for proton transfer and is gated by conformational changes in several TMHs, including ND1-TMH4 and ND6-TMH3 (Figure 6A).

**Figure 6.**
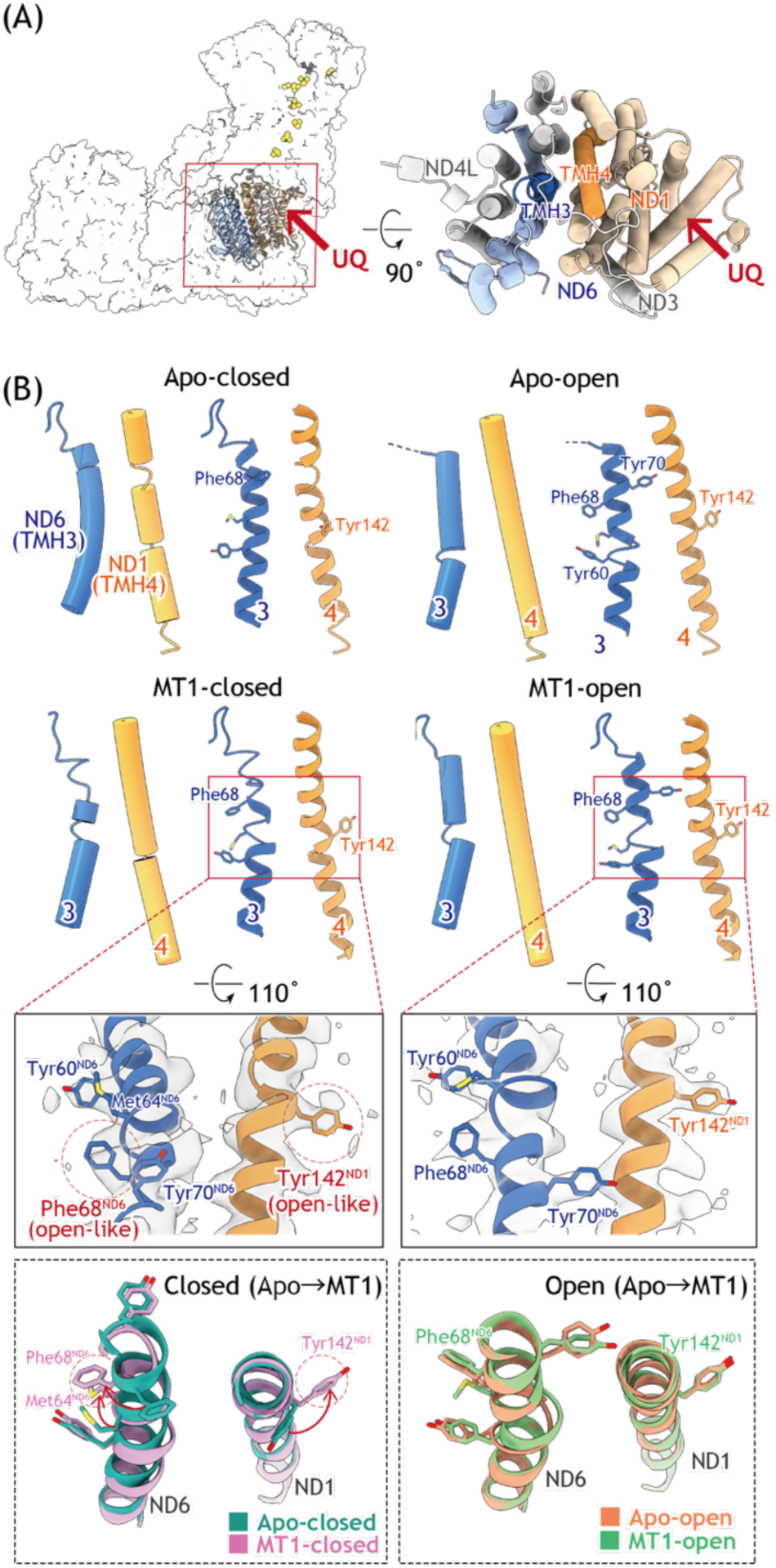
MT1 induces open-like local arrangement of ND6-TMH3 and ND1-TMH4 in both closed and open conformations of bovine complex. **I.** (A) Location of ND6-TMH3 (*blue*) and ND1-TMH4 (*orange*) surrounding the entrance of the UQ-accessing tunnel. The boxed region in the overall structure (*left*) is enlarged and rotated by 90° (*right*). The *red arrows* indicate the entrance of the UQ-accessing tunnel. (B) Structural comparison of ND6-TMH3 and ND1-TMH4 regions in the apo and MT1-bound structures. The *upper panels* show cylinder and ribbon representations of ND6-TMH3 and ND1-TMH4 in the apo-*closed (left)*- and apo-*open (right)-*conformations. The *middle panels* show the corresponding regions in the MT1-bound *open (left)*-and *closed (right)*-conformations, with cryo-EM densities displayed around the key residues Tyr142 of ND1 and Phe68 of ND6. In the MT1-bound closed conformation, Tyr142 of ND1 and Phe68 of ND6 adopt *open-like*-conformations resembling those observed in the apo-*open* structure (red dashed circles). In contrast, these residues undergo little additional rearrangement upon MT1 binding in the *open conformation*. The *lower panels* compare the apo and MT1-bound structures by superposition for the *closed (left)*- and *open (right)*-conformations, highlighting the MT1-induced *open-like* local arrangement of ND6-TMH3 and ND1-TMH4.

In the closed conformation, ND1-TMH4 adopts a bent conformation, whereas the adjacent ND6-TMH3 is α-helical throughout, with the side chain of ND6-Phe68 oriented toward ND1 (Figure 6B). This arrangement is considered to support formation of a continuous water-mediated network extending from the UQ-accessing tunnel toward membrane domain [*11–13*]. In the open conformation, straightening of ND1-TMH4 is accompanied by repositioning of ND1-Tyr142, whereas ND6-TMH3 develops a π-bulge that results in an approximately 180° rotation of the matrix-side segment of the helix containing ND6-Phe68 (Figure 6B). These conformational rearrangements are hypothesized to disrupt the continuity of the water-mediated network, which is critical for redox-driven proton translocation.

In contrast to the PSST loops, MT1-bound complex I retained the open state orientation of ND1-Tyr142, ND6-Met64, and ND6-Phe68 in both open and closed conformations (Figure 6B). In the ND1 subunit, clear densities of Tyr142 were observed in both MT1-bound conformations, showing that the rotation of Tyr142 relieved the distortion of TMH4 (Figure 6B). In the ND6 subunit, although the density corresponding to ND6-Phe68 was relatively weak in the MT1-bound closed conformation, TMH3 clearly retained a π-bulge-like conformation in both MT1-bound structures, indicating that mycothiazole stabilizes the open state arrangement of ND6-TMH3 (Figure 6B). Together with the rotation of ND1-Tyr142, these observations indicate that MT1 binding induces coordinated rearrangements of ND1-TMH4 and ND6-TMH3 toward their open state-like arrangement in the closed conformation. Thus, despite their different global conformations, the MT1-bound open and closed structures converge on a common hybrid arrangement around the UQ-accessing tunnel; namely, closed-conformation-like PSST loops and open-conformation-like ND1-TMH4 and ND6-TMH3.

### Bound mycothiazole in UQ-accessing tunnel

In both open and closed conformations, we observed two distinct densities within the UQ-accessing tunnel (Figure 7). One density was observed in the deep region of the tunnel, and the other was located near the entrance of the tunnel formed by ND1. The density in the deep region is surrounded by the β1– β2 strands of the 49 kDa subunit and two loops of the PSST subunit, both of which underwent MT1-induced conformational rearrangements (Figure 7A). The β1–β2 strands overlap with the region identified by photoaffinity labeling using [¹²⁵I]pMT (Figure 3D). Together, these observations support the interpretation that the major density in the deep region corresponds to bound MT1. However, the local resolution was insufficient to unambiguously define the inhibitor orientation or specific interactions with key residues implicated in the binding and redox chemistry of UQ (Tyr108^49k^, His59^49k^, and Asp160^49k^, Figure 7B) [*17,49,50*].

**Figure 7.**
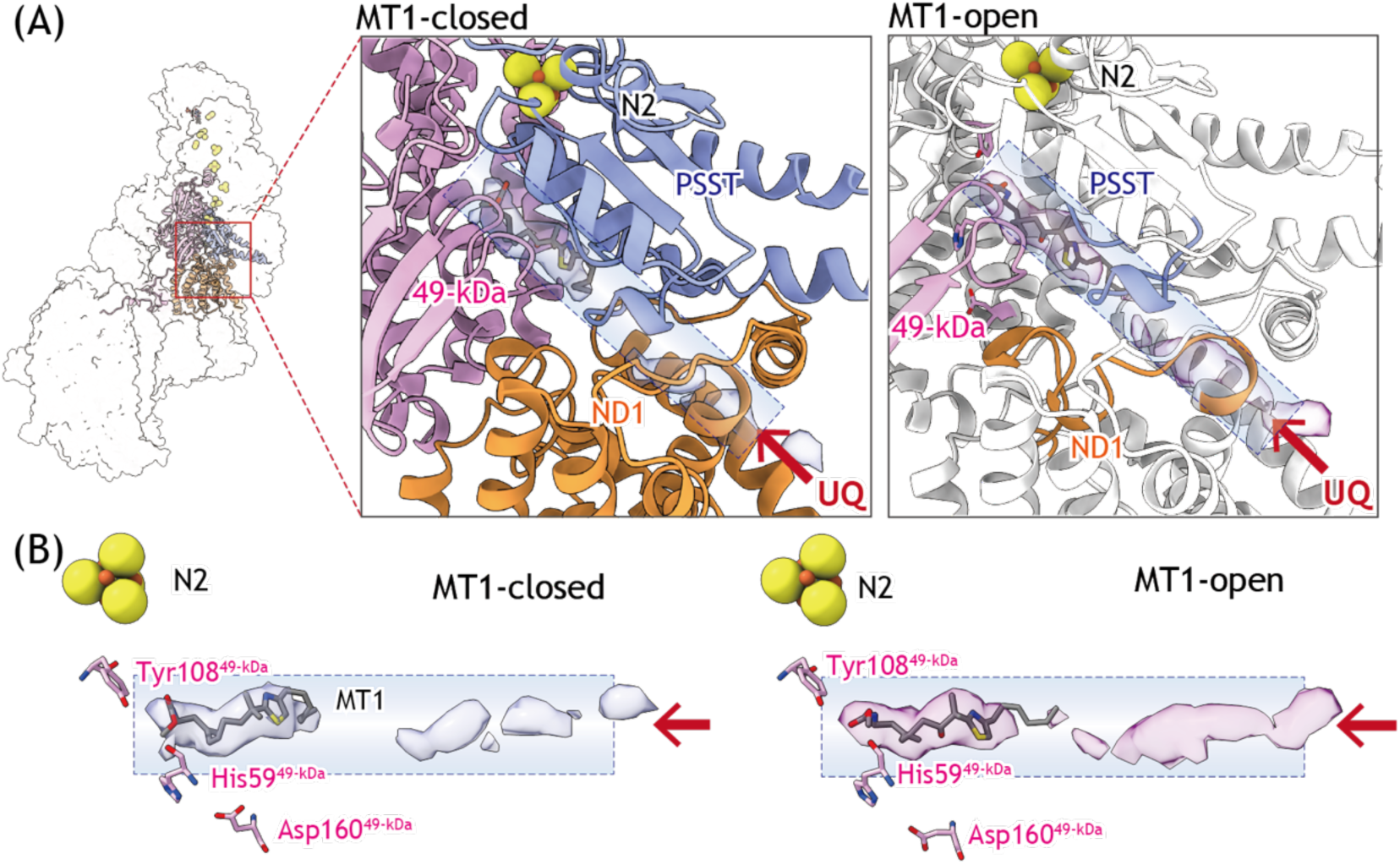
Proposed MT1-binding site within UQ-accessing tunnel of bovine complex. **I.** (A) Overall view of the unmodeled cryo-EM density within the UQ-accessing tunnel in *closed (left)-* and *open (right)*-conformations of bovine complex I. The unmodeled cryo-EM density assigned to MT1, shown in *transparent light blue* and *light pink* for the *closed*- and *open conformation*s, respectively, is located in the deep region of the UQ-accessing tunnel and surrounded by β1–β2 strands of the 49 kDa subunit (*magenta*), PSST subunit (*blue*), and ND1 subunit (*orange*). In the *open conformation*, only the regions that became sufficiently ordered to be modeled upon MT1 binding are highlighted in *color*, whereas the regions already modeled in the apo-structure are shown in *white*. The *red arrow* indicates the entrance of the UQ-accessing tunnel. (B) Close-up views of the density in the *closed (left)-* and *open (right)*- conformations. The density is shown as *transparent surfaces in light blue* and *light pink* for the *closed*- and *open conformation*s, respectively, together with key residues involved in UQ recognition: His59, Tyr108, and Asp160 of the 49 kDa subunit. The local resolution was insufficient to unambiguously determine the orientation of MT1 or its specific interactions with these key amino acid residues.

The weak density near the tunnel entrance is located close to the ND1 region labeled by [^125^I]pMT (Figure 2C). Excess IACS-010759, which binds to the ND1 subunit [*32*], suppressed the labeling of ND1 by [^125^I]pMT (Figure 2B). These results suggest that the binding regions of mycothiazole and IACS-010759 partially overlap in ND1. Therefore, it is likely that the minor density near the tunnel entrance corresponds to bound MT1.

Nevertheless, this assignment of the densities requires caution because weak, discontinuous unassigned densities were also present at several positions along the UQ-accessing tunnel in our previously reported inhibitor-free closed structure (Figure S5) [*48*]. Direct comparison with the inhibitor-free open conformation is limited because several elements forming the tunnel were unresolved in that structure. In contrast, the structures obtained in the presence of MT1 exhibit more continuous densities, particularly in the deep region of the tunnel. Although contributions from endogenous lipids or UQ cannot be excluded, the locations of densities, together with the photoaffinity-labeling results and MT1-induced ordering of the surrounding structural elements, support the interpretation that MT1 contributes substantially to the observed densities. Accordingly, this means that two molecules of MT1 can occupy the interior of the UQ-accessing tunnel. Multiple inhibitor occupancy within the tunnel is not unprecedented, being reported for rotenone [*12*] and piericidin A [*23*]. However, because a high concentration of MT1 (500 µM) was used for cryo-grid preparation, the second density near the entrance may reflect an artifact caused by the experimental conditions, as pointed out for cryo-EM works by *Kampjut et al.* [*12*].

The ND5 subunit was also labeled by [¹²⁵I]pMT (Figures 3A and 3B), whereas no density suggestive of bound MT1 or clear local structural rearrangement was observed around the ND5 body at the distal end of the membrane domain in either conformation. Although the significance of ND5 labeling remains unclear, previous studies indicated that there may be a hydrophobic cleft around the long lateral helix of ND5, which extends alongside ND4 and ND2, and the cleft accommodates hydrophobic chemicals such as UQ derivatives and rotenone [*12,44*]. Notably, numerous phospholipid molecules are bound in and around this region in the present structures (Figure S6), indicating a highly hydrophobic local environment. Such a lipid-rich environment may facilitate the transient association of [^125^I]pMT with this region and thereby contribute to the ND5 labeling observed. Nevertheless, the present cryo-EM study provided no clear evidence for the existence of bound MT1 in this region.

## Discussion

### Mycothiazole reveals architecture of UQ-accessing tunnel in open conformation

A hallmark of the open conformation of mammalian complex I is the high-level flexibility of the four loop regions surrounding the UQ-accessing tunnel: the loop connecting β1–β2 strands of the 49 kDa subunit, that between TMHs 5 and 6 of ND1, that between TMHs 1 and 2 of ND3, and that between TMHs 4 and 5 of ND6 [*12*]. A notable finding of the present cryo-EM study was that the binding of mycothiazole results in a distinct conformation of these loops, enabling structural characterization of the whole UQ-accessing tunnel in the open conformation of bovine complex I. Previous cryo-EM studies also reported inhibitor-bound open conformations of complex I, including rotenone-bound [*12*] and proguanil-bound (hydrophobic analogue of biguanide-type inhibitor [*28*]) complex I. However, these open conformation structures retained substantial disorders in the matrix side of the UQ-accessing tunnel. For example, in the rotenone-bound open state, the binding of rotenone induced ordering of the 49 kDa and PSST loops, whereas the ND1 and ND3 loops remained disorders [*12*]. Similarly, proguanil-bound open structures also retained the characteristic disorder associated with the open conformation [*28*]. In contrast, the binding of mycothiazole gave rise to a distinct conformation of these structural elements (Figure 4) and, consequently, we could model the entire UQ-accessing tunnel. The regions in the 49-kDa and ND1 subunits, that were stabilized in the presence of MT1, correspond to the binding region of [^125^I]pMT identified by photoaffinity labeling experiments (Figures 3 and 4). Thus, the present study provides the first entire structural view of the UQ-accessing tunnel in the “open” conformation of mammalian complex I.

### Mycothiazole-bound open conformation is distinct from biochemically defined deactive state of complex I

As the open conformation has often been considered as the structural counterpart of the biochemically defined “deactive” state of complex I, it is important to distinguish these two structures. The open and closed conformations were originally defined on the basis of cryo-EM analyses [*11,12,51*], whereas the active and deactive states were originally defined on the basis of biochemical properties of the enzyme [*52,53*]. In particular, the deactive state is characterized by accessibility of thiol-reactive reagents to the conserved Cys39 located in the ND3 TMH1–2 loop [*53*]. Although similarities between the open conformation and deactive state have been noted in several studies because both involve marked rearrangements of the flexible loop structures around UQ-accessing tunnel [*11,51*], the precise relationship between these structural conformations and biochemical states remains to be elucidated.

The mycothiazole-bound open conformation identified in the present study does not fully satisfy the criteria established for the biochemically defined deactive state. In particular, the conformation of the ND3 TMH1–2 loop observed in the MT1-bound open conformation is considered to be incompatible with the accessibility of Cys39 against externally added thiol-reactive reagents (Figure S4). Moreover, the four structural elements surrounding the UQ-accessing tunnel above, which became fully ordered in the presence of mycothiazole, adopted conformations closely resembling those observed in the closed conformation (Figures 4 and S4). Collectively, these observations suggest that the mycothiazole-bound open conformation is unlikely to correspond to the biochemically defined deactive state. Rather, mycothiazole may shift part of the UQ-catalytic region toward a closed state-like architecture while preserving the overall open conformation of the UQ-accessing tunnel. These findings indicate that the mycothiazole-bound open conformation is distinct from the biochemically defined deactive state, and support the binding of a variety of ligand molecules (including substrate UQs) within the open conformation of mammalian complex I [*29,30,32–34,45,54*].

### Mycothiazole induces hybrid structural state around UQ-catalytic site

The functional significance of the open conformation is a matter of ongoing debate since premature protonation of reduced UQ intermediates at the “top” of the tunnel from the matrix side will result in loss of energy [*12,51*]. Some studies have associated the open conformation with the biochemically defined deactive state [*51*], whereas others have proposed that it participates in catalytically relevant structural transition during the enzyme turnover [*12*]. Although the present study provides no direct answer to this question, a combination of unusual structural features around the UQ-catalytic site hitherto unknown was revealed. Namely, mycothiazole binding does not alter the global architecture of complex I in either open or closed conformations, whereas it specifically rearranges local structural elements surrounding the UQ-accessing tunnel. In the open conformation, mycothiazole shifts the PSST loops toward a closed-conformation-like arrangement, whereas ND1-TMH4 and ND6-TMH3 retain their open-conformation-like configurations (Figures 5 and 6). Conversely, in the closed conformation, the PSST loops retain their closed-conformation-like arrangement, whereas ND1-TMH4 and ND6-TMH3 shift toward open-conformation-like configurations (Figures 5 and 6).

Thus, despite their different global conformations, the MT1-bound open and closed structures converge toward a common “hybrid” arrangement, in which closed-conformation-like PSST loops coexist with open-conformation-like ND1 and ND6 segments. These observations suggest that the conformational landscape of mammalian complex I is more complex than a simple *open-versus-closed* structural classification. The structural elements surrounding the UQ-accessing tunnel may respond independently to the binding of a variety of ligand molecules, generating hybrid conformations. In particular, local structural plasticity of the matrix-side interfacial region of the UQ-accessing tunnel is evident. For example, Masuya et al. [*34*] demonstrated that 49-kDa Asp160 located deep inside the tunnel and ND3 Cys39 on the matrix-side flexible TMH1-2 loop can be directly linked by bulky bifunctional chemical cross-linkers. We also showed that bulky UQ analogues and membrane-anchored quinazoline inhibitors, both unlikely to access the catalytic site through the canonical UQ-accessing tunnel, can nevertheless react with or inhibit complex I [*55,56*]. Overall, the present findings further highlight the pronounced conformational plasticity of the UQ-accessing tunnel in mammalian complex I.

The local structural rearrangement induced by ligand binding may have functional consequences related to UQ redox chemistry in complex I, such as generation of reactive oxygen species [*57*]. In fact, structurally diverse bioactive reagents targeting complex I (e.g., IACS-010759 [*31*], S1QELs [*30,57*], and mubritinib [*58*]) have been identified through intensive screening of large compound libraries. Given the diverse action manners of ligand molecules targeting complex I, there may be considerable room to further develop unique reagents that exhibit a broad spectrum of biological activities [*59*]

## Materials and methods

### Synthesis of compounds used in this study

The syntheses of MT1, pMT, and [^125^I]pMT are summarized in Schemes S1−S3, and the detailed procedures are described in supporting information. Other inhibitors used in this study were the same sample as those used previously [*32,45*].

### Preparation of bovine heart SMPs

SMPs were prepared from bovine heart mitochondria according to the procedures described by Matsuno-Yagi and Hatefi [*60*], and stored in a buffer containing 250 mM sucrose and 10 mM Tris/HCl (pH 7.4) at -80°C until use. NADH-UQ_1_ oxidoreductase activity in SMPs was measured spectrophotometrically by following the oxidation of NADH with a Shimadzu UV2600i instrument (340 nm, ε = 6.2 mM^-1^cm^-1^) at 30°C. The reaction medium (2.5 mL) contained 250 mM sucrose, 1.0 mM MgCl_2_, 0.2 µM antimycin A, 4.0 mM KCN, 50 µM UQ_1_, and 50 mM potassium phosphate buffer (pH 7.4). The final protein concentration was set to 30 µg of protein/mL. The reaction was initiated by the addition of 50 µM NADH after the equilibration of SMPs with inhibitor for 4 min. IC_50_ values of the inhibitors were estimated by Prism 11 (GraphPad).

### Photoaffinity labeling of complex I in SMPs

For photoaffinity labeling by [^125^I]pMT and [^125^I]AzQ, bovine SMPs (2.0 mg of protein/mL) were incubated with [^125^I]pMT (2.0 nM) or [^125^I]AzQ (5.0 nM) in buffer containing 250 mM sucrose, 5.0 mM MgCl_2_, and 50 mM KPi (pH7.4) at room temperature for 5 min [*32,41,42*]. After the addition of 50 µM of NADH, the mixture was irradiated with a long wavelength UV lamp (Black-ray model B-100A, UVP) on ice for 10 min at a distance of 10 cm from the light source. When the competition test was conducted, SMPs were incubated with different types of complex I inhibitors for 10 min at room temperature prior to treatment with [^125^I]pMT.

SMPs labeled by [^125^I]pMT were solubilized in sample buffer containing 1% (w/v) *n*-dodecyl-β-D-maltoside, 0.75 M aminocaproic acid, and 50 mM Bis-Tris/HCl (pH 7.0) on ice for 1 h, and the labeled complex I was isolated by BN-PAGE using a hand-cast 6% isocratic gel [*32*]. Then, 50 mM Tricine and 15 mM Bis-Tris/HCl (pH 7.0) and 50 mM Tris/HCl (pH 7.0) were used as cathode and anode electrode buffers, respectively. The subunits of labeled complex I were separated on a 12.5% Laemmli gel [*61*] or 10% Schägger-type gel (10% T, 3% C, containing or not containing 6.0 M urea) [*62*], then stained with CBB or silver (Wako Silver stain MS kit, Wako Pure Chemicals), dried, exposed to an imaging plate (BAS-MS2040, Fuji Film), and visualized with the bio-imaging analyzer Typhoon-FLA 9500 (Cytiva). The incorporated radioactivity of each band was quantified using Image Quant (Cytiva).

Doubled SDS-PAGE was conducted as described previously [*32,47*]. In brief, the labeled complex I was separated on a first-dimensional 10% Schägger-type gel (10% T, 3% C, containing 6.0 M urea). The gel slice was then acidified with 100 mM Tris-HCl (pH 2.0) for 30 min, followed by second-dimensional separation on a 16% Schägger-type gel (16% T, 3% C) [*62*].

### Protein analyses

Regarding the partial digestion (Cleveland mapping) of labeled subunits, CBB-stained bands of the target subunits were digested with Glu-C (Roche Applied Science) in a 20% Tris-EDTA mapping gel according to previously described procedures [*41,47*].

To exhaustively digest the labeled subunits, target subunits were recovered from the SDS gel by electroelution or direct diffusion in 10 mM Tris-HCl buffer (pH 8.0) containing 0.025% (w/v) SDS. The isolated subunits were digested with lysyl endopeptidase (Wako Pure Chemicals) or endoprotease Asp-N (Roche Applied Science) in 50 mM Tris HCl-buffer containing 0.1% SDS or 50 mM NaPi buffer containing 0.01% SDS, respectively. The digests were resolved on a Schägger-type SDS gel [*62*], dried, and then exposed to an imaging plate (BAS-MS2040, Fuji Film).

### Cryo-EM sample preparation and data collection

SMPs (6.5 mg/mL) were mixed with mycothiazole to a final concentration of 500 μM and incubated for 60 min at 4°C. Then, 2.7 μL of the sample was applied to a glow-discharging Quantifoil UltraAufoil R1.2/1.3 grid using Vitrobot Mark IV (Thermo Fisher Scientific). The grid was blotted for 3 s with a blot force of 5 at 4 °C and 100% humidity, and then rapidly frozen in liquid ethane. Cryo-EM movies were automatically acquired using a Titan Krios G2 (Thermo Fisher Scientific) equipped with a K3 electron detector (Gatan) using SerialEM software [*63*]. The movies were collected at a nominal magnification of 29,000 and pixel size of 0.84 Å/pix. The total electron dose was 50 electrons/Å^2^, and the defocus range was -0.8 to -1.8 μm.

### Image processing

Image-processing steps are summarized in Figure S2A. All steps were performed using CryoSPARC v4.7.0 [*64*]. Topaz software was applied for machine-learning-based particle picking [*65*]. The acquired movies were corrected with Patch Motion Correction. Contrast transfer function (CTF) was estimated using Patch CTF estimation. Particles were picked with Topaz and extracted at 1.20 Å/pixel. Extracted particles were subjected to 2D classification, heterogeneous refinement, and 3D classification to remove junk particles. The selected particles were subjected to homogeneous refinement for reconstruction of the structure of supercomplexes (SCs). To obtain high precision CTF parameters, the 201,744 particles of SCs were subjected to local refinement in the complex III region, followed by global/local CTF refinement. To classify the complex I state, heterogeneous refinement was performed on particles with CTF parameters refined in the complex III region, and selected particles were subjected to local refinement in complex I. These procedures provided density maps for *open* MT1 and *closed*-MT1 at overall resolutions of 3.4 and 3.5 Å, respectively. Resolution was estimated by CryoSPARC v4.7.0 using the gold-standard Fourier shell correlation of 0.143.

### Model building and refinement

The atomic models for *open*-MT1 and *closed*-MT1 were built based on our previously determined cryo-EM structures of bovine complex I in SMPs (PDB: 9W2U and 9W2V). The initial model was fitted into the density map by ChimeraX [*66*] and refined using the phenix_real_space_refine program [*67*]. The resulting model was manually corrected by Coot [*68*] and ISOLDE [*69*] and refined using phenix.real-space refinement until model parameters improved. The model quality was evaluated with MolProbity [*70*]. The statistics of maps and models presented here are summarized in Tables S1. All figures were prepared using ChimeraX.

### Data availability

All study data are included in the article and/or in *Supporting Information*. The cryo-EM maps have been deposited in the EMDB under accession codes, EMD-81928 (MT1-closed) and EMD-81929 (MT1-open). The atomic models have been deposited in the Protein Data bank under accession codes, 43JM (MT1-closed) and 43JN (MT1-open).

## Supporting information

Figures S1-S6, Table S1, and Synthetic procedures

## Acknowledgments and funding sources

We thank the Radioisotope Research Center, Kyoto University, for technical support in the radioisotope experiments. We acknowledge the Research Support Project for Life Science and Drug Discovery (Basis for Supporting Innovative Drug Discovery and Life Science Research (BINDS)) from AMED under Grant Number JP17am0101001 (support number 1312). This work was financially supported by JSPS KAKENHI (24K08729 to T.M., 21H02130 to H.M., and 25K01958 to M.M.), the Uehara Memorial Foundation (to M.M.), and Tokyo Kasei Chemical Promotion foundation (to M.M.).

