## Supplementary material for "Mycothiazole enables structural characterization of entire ubiquinone-accessing tunnel in mitochondrial respiratory complex I": Figures S1-S6, Table S1, and Synthetic procedures

*[1]Deparment of Molecular Biosciences, Kyoto Sangyo University, Kyoto, Japan, [2]Research and Development, Sumitomo Pharma. Co., Ltd., Osaka, Japan, [3]Division of Applied Life Sciences, Graduate School of Agriculture, Kyoto University, Kyoto, Japan, [4]Department of Applied Biological Chemistry, Graduate School of Agriculture, Osaka Metropolitan University, Osaka, Japan, [5]Research Center for Ultra-High Voltage Electron Microscopy, Osaka University, Osaka, Japan*

**Figure S1:** Summary of the binding site of [$^{125}$I]AzQ in complex I.

**Figure S2:** Summary of the binding site of [$^{125}$I]pMT in complex I.

**Figure S3:** Cryo-EM data-processing workflow and resolution assessment of bovine complex I.

**Figure S4:** Comparison of MT1 bound open- and closed-structures of the ND1, 49 kDa, ND3 and ND6 subunits.

**Figure S5:** Cryo-EM density corresponding to the UQ-accessing tunnel in apo-complex I in the closed conformation.

**Figure S6:** Lipid-rich environment surrounding the long lateral helix of ND5.

**Table S1:** Statistics for Cryo-EM data, refinement, and validation of complex I with bound mycothiazole.

**Synthetic procedures of MT1, pMT, and [$^{125}$I]pMT**

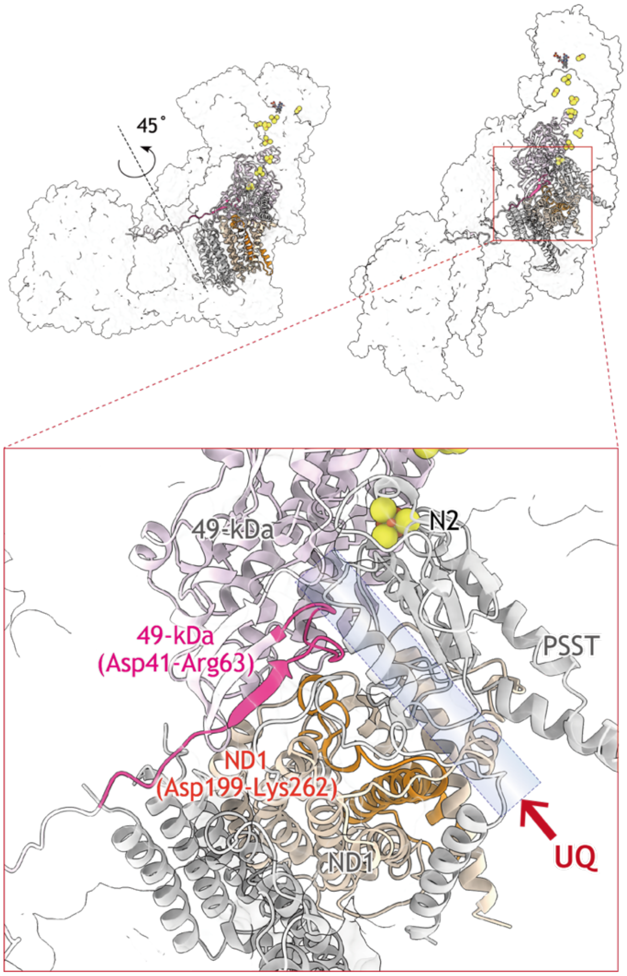


**Figure S1. Summary of the binding sites of [$^{125}$I]AzQ in complex I.** The upper panels show two views of complex I related by a 45° rotation, and the lower panel shows a close-up view of the boxed region. The regions labeled by the photoreactive quinazoline ([$^{125}$I]AzQ) in the 49-kDa (Asp41–Arg63) and ND1 (Asp199–Lys262) subunits are shown in *magenta* and *orange*, respectively. The putative UQ-accessing tunnel is indicated by the transparent blue region, and the entrance to the tunnel is indicated by a *red arrow*. The MT1-bound closed conformation of bovine complex I was used as the structural template (PDB entry 43JM).

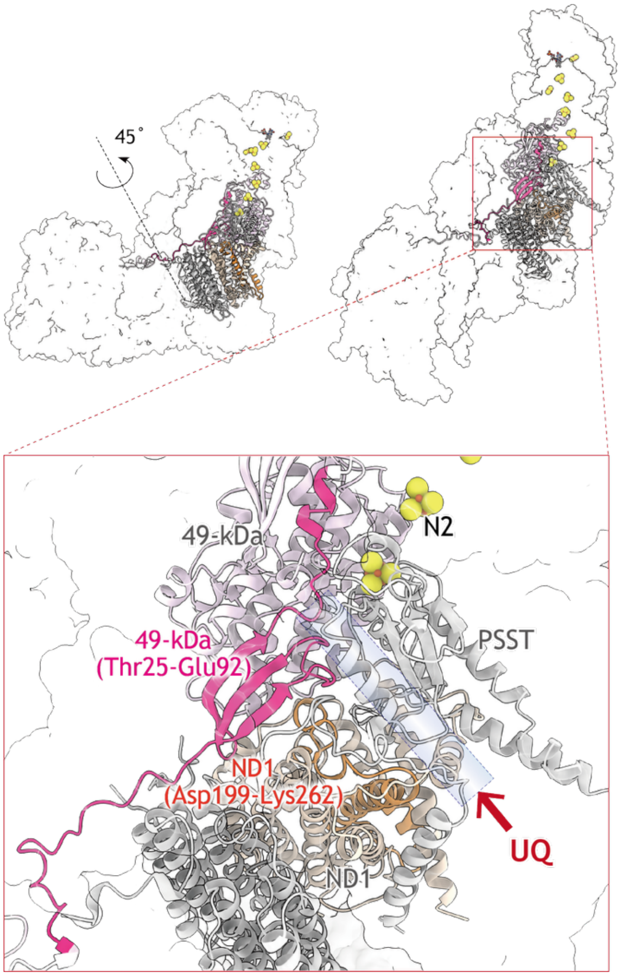


**Figure S2: Summary of the binding sites of [$^{125}$I]pMT in complex I.** The upper panels show two views of complex I related by a 45° rotation, and the lower panel shows a close-up view of the boxed region. The regions labeled by the photoreactive mycothiazole ([$^{125}$I]pMT) in the 49-kDa (Thr25–Glu92) and ND1 (Asp199–Lys262) subunits are shown in *magenta* and *orange*, respectively. The putative UQ-accessing tunnel is indicated by the transparent blue region, and the entrance to the tunnel is indicated by a *red arrow*. The MT1-bound closed conformation of bovine complex I was used as the structural template (PDB entry 43JM).

(A)

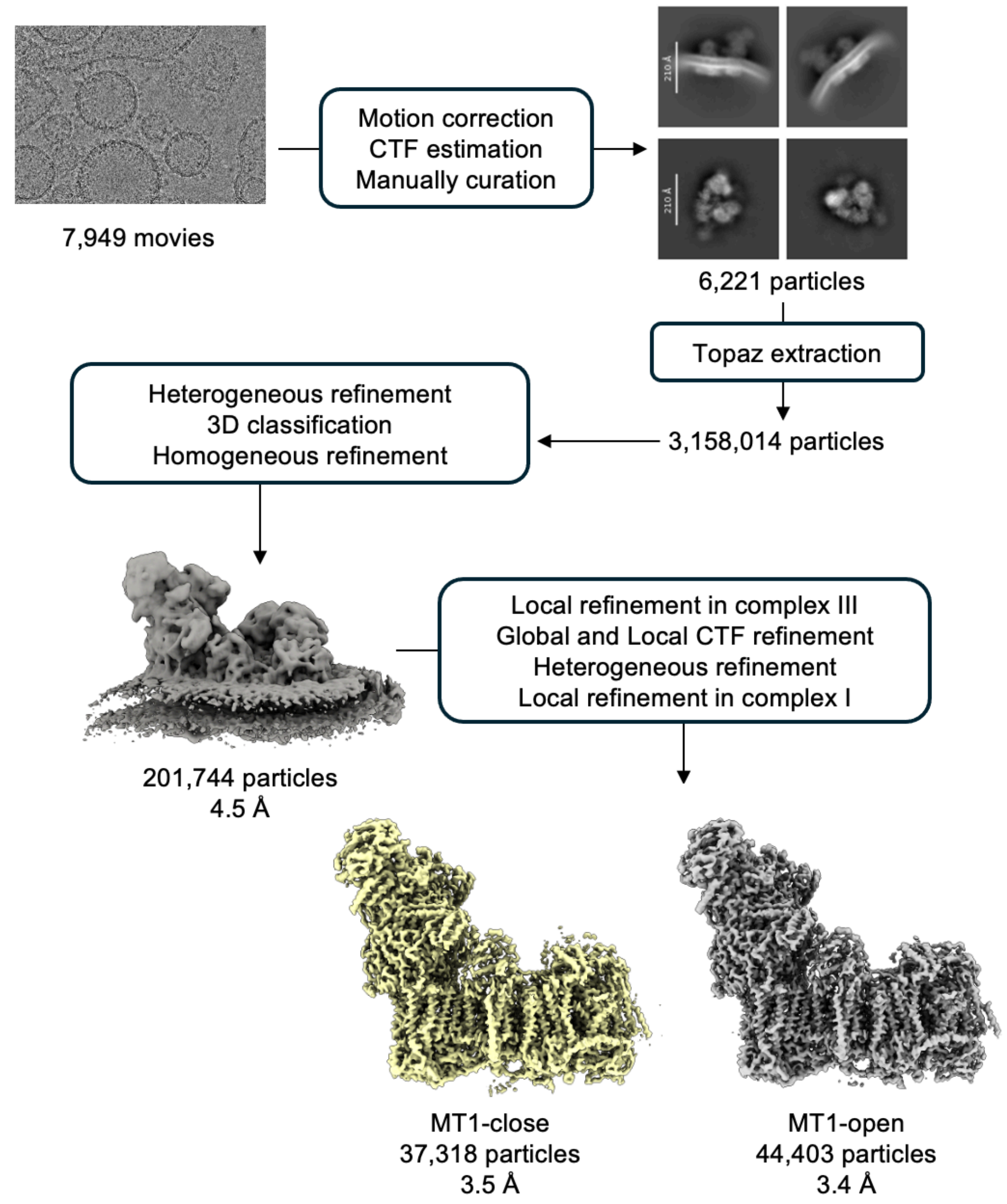


**Figure S3 (continued)**

(B)

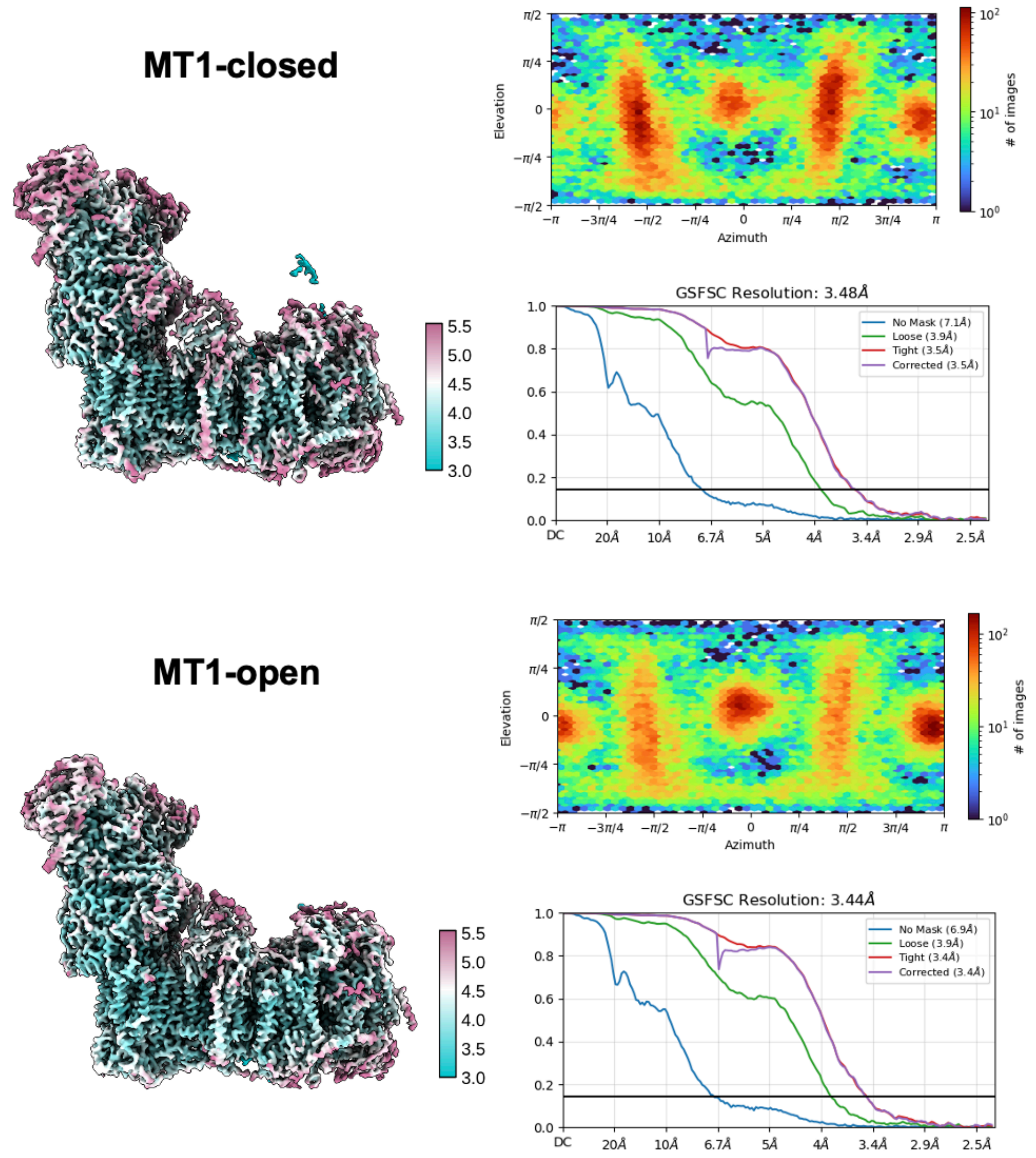


**Figure S3. Cryo-EM data-processing workflow and resolution assessment of bovine complex I.** (A) Workflow for single-particle cryo-EM analysis using cryoSPARC. (B) Resolution assessment of the reconstructed maps. The *left panels* show the local-resolution maps, the *upper right panels* show the angular distributions of particles, and the *lower right panels* show the gold-standard Fourier shell correlation (GSFSC) curves.

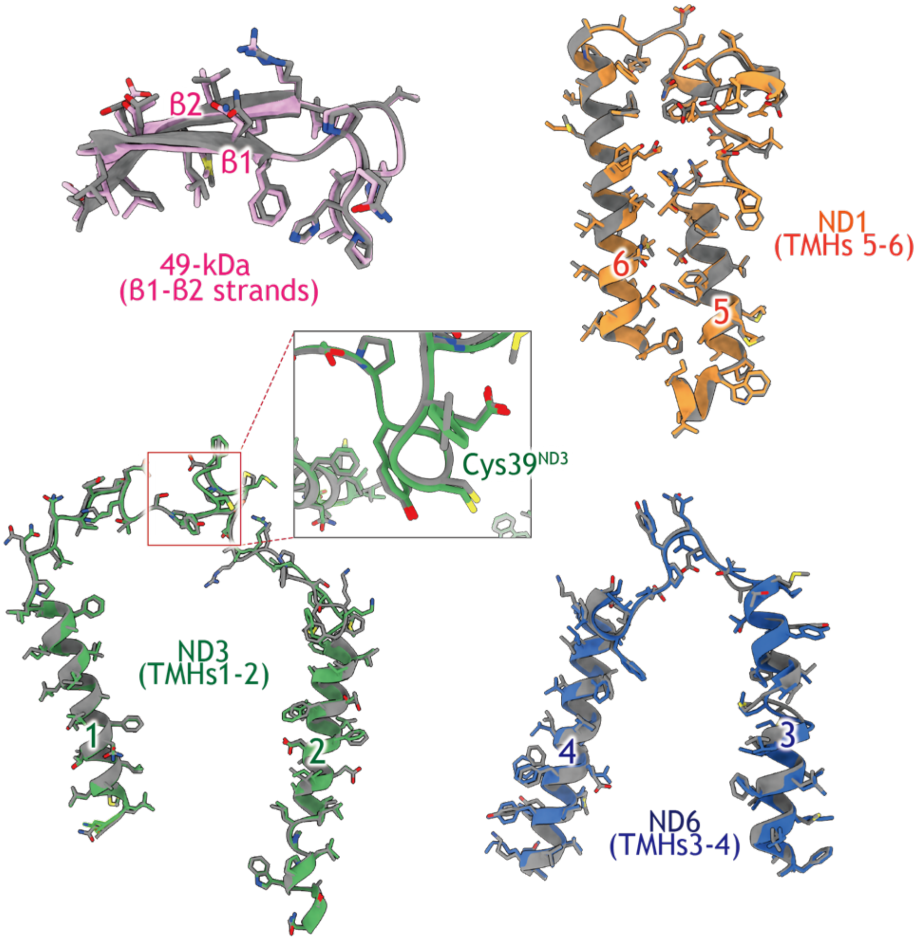


**Figure S4: Comparison of MT1-bound complex I in the open and closed conformations.** The *gray* and *colored* models represent the MT1-bound closed and open conformations, respectively. Structural comparisons are shown for the β1–β2 strands of the 49-kDa subunit, TMHs 5–6 of ND1, TMHs 1–2 of ND3, and TMHs 3–4 of ND6. The inset highlights the region surrounding Cys39 of ND3.

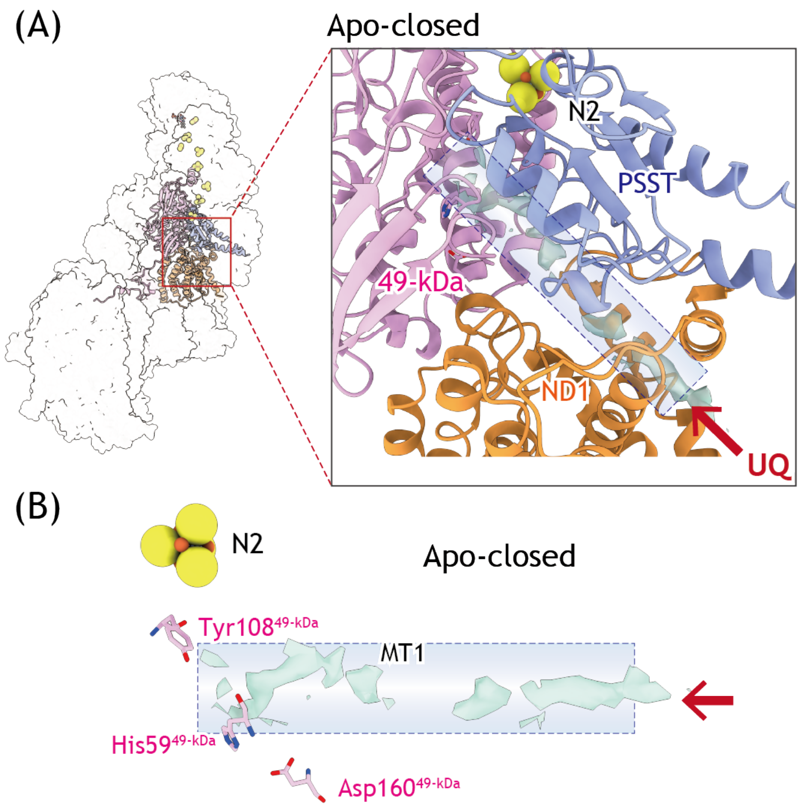


**Figure S5: Cryo-EM density corresponding to the UQ-accessing tunnel in apo-complex I in the closed conformation.** (A) Unmodeled cryo-EM density observed within the UQ-accessing tunnel of apo-complex I in the closed conformation (PDB entry: 9W2V) is shown in *transparent light green*. The 49-kDa, PSST, and ND1 subunits, which form the UQ-accessing tunnel, are shown in *magenta*, *blue*, and *orange*, respectively. (B) Close-up view of the unmodeled density, shown in *light green*, together with key residues involved in UQ recognition: His59, Tyr108, and Asp160 of the 49-kDa subunit.

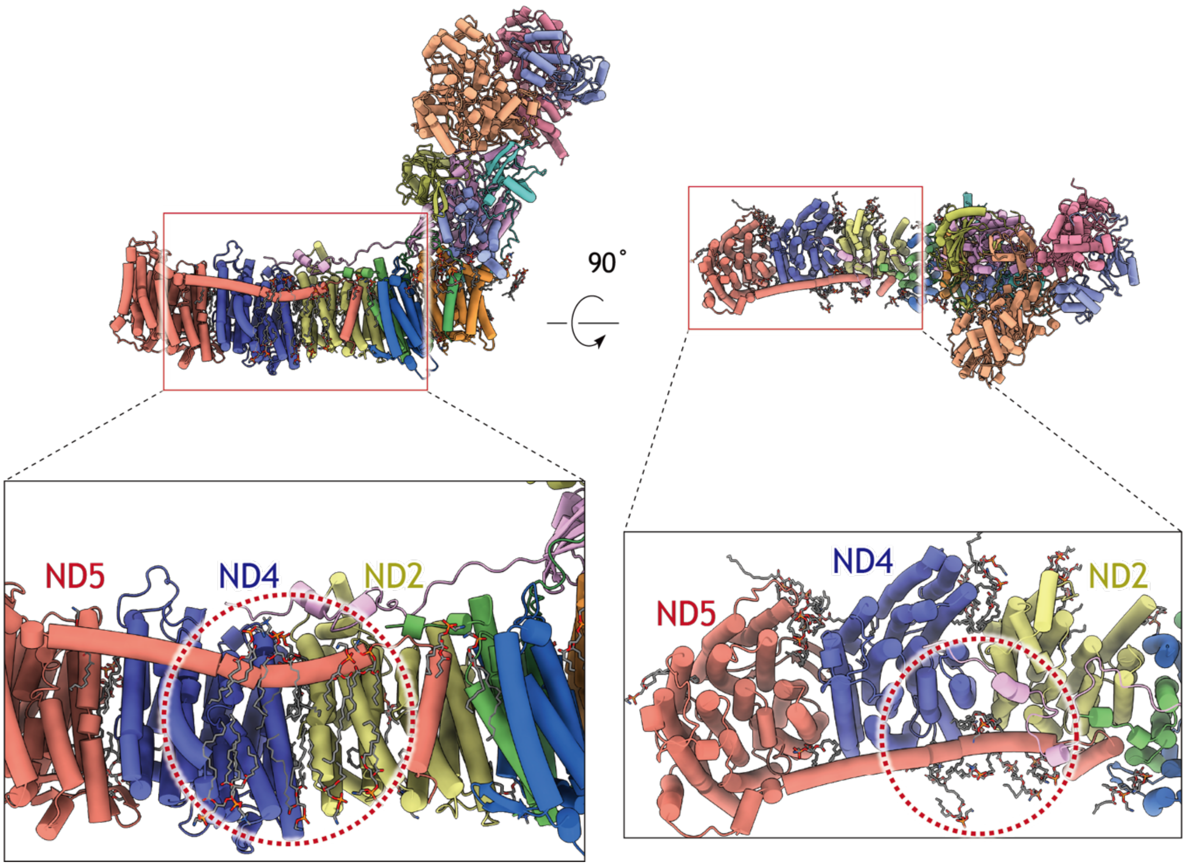


**Figure S6. Lipid-rich environment surrounding the long lateral helix of ND5.** The MT1-bound closed conformation of bovine complex I (PDB entry 43JM, only 14 core subunits are presented) is shown in two orientations related by an approximately 90° rotation. The boxed regions are enlarged in the lower panels. ND5, ND4, and ND2 are shown in *salmon*, *blue*, and *yellow*, respectively, and modeled phospholipid molecules are shown as sticks. The *dashed circles* indicate the hydrophobic region around the long lateral helix of ND5 at the interface with ND4 and ND2. Numerous phospholipid molecules are modeled in and around this region, whereas no additional density attributable to bound MT1 was identified.

**Table S1: Statistics for Cryo-EM data, refinement, and validation of complex I with bound mycothiazole**

| | **MT1-closed** | **MT1-open** |
|---|---|---|
| EMDB ID | EMD-81928 | EMD-81929 |
| PDB ID | 43JM | 43JN |
| **Data collection and processing** | | |
| Magnification | 29,000 | 29,000 |
| Voltage (kV) | 300 | 300 |
| Total dose ($e^{-}/Å^{2}$) | 50 | 50 |
| Defocus range (µm) | -0.8 to -1.8 | -0.8 to -1.8 |
| Pixel size (Å) | 0.84 | 0.84 |
| Number of initial particles | 3,158,014 | 3,158,014 |
| Number of final particles | 37,318 | 44,403 |
| Map resolution (Å) | 3.48 | 3.44 |
| FSC threshold | 0.143 | 0.143 |
| Symmetry imposed | C1 | C1 |
| Map sharpening B-factor ($Å^{2}$) | -41.4 | -47.3 |
| **Model Refinement** | | |
| Initial model used | 9W2V | 9W2U |
| Model resolution (Å) | 3.8 | 3.8 |
| FSC threshold | 0.5 | 0.5 |
| **Composition** | | |
| Nonhydrogen atoms | 68,647 | 68,724 |
| Residues | 8,288 | 8,286 |
| Ligands | GTP: 1, MG:1, K:1, SF4:6, FMN:1, CDL:6, ZN:1, FES:2, PC1:12, NDP:1, A1MID (mycothiazole):1, 3PE:19 | GTP: 1, MG:1, CHD:1, K:1, SF4:6, MYR:1, FMN:1, CDL:7, ZN:1, FES:2, PC1:13, NDP:1, A1MID (mycothiazole):1, 3PE:18 |
| Bonds length (Å) | 0.006 | 0.007 |
| Bonds Angles (° ) | 1.254 | 1.354 |
| MolProbity score | 1.86 | 1.94 |
| Clash score | 8.26 | 8.16 |
| Rotamer outliers (%) | 2.32 | 2.66 |
| **Ramachandran plot** | | |
| Favored (%) | 97.28 | 96.98 |
| Allowed (%) | 2.69 | 2.96 |
| Outliers (%) | 0.04 | 0.06 |

### Synthetic procedures of MT1, pMT, and [$^{125}$I]pMT

#### *General procedures*

All moisture- and air-sensitive reactions were performed in oven-dried glassware under a nitrogen (or argon) atmosphere with dry solvents using standard syringe septum techniques. $^{1}$H-NMR spectra were recorded at 400 MHz or 500 MHz with Bruker AVANCE III 400 or AVANCE III 500 spectrometers, respectively, using tetramethylsilane (TMS) as an internal standard. $^{13}$C-NMR spectra were recorded at 100 MHz or 125 MHz. Chemical shifts ($\delta$) are given in ppm relative to TMS with coupling constants ($J$) in Hz. The mass spectra were recorded on a Shimadzu LCMS-8040 with ESI source. Thin-layer chromatography (TLC) was performed on Merk TLC plate silica-gel 60F$^{254}$, and the spot was detected by iodine, anis, phosphomolybdic acid or UV absorbance. Dry solvents were used either as purchased or freshly distilled using common practices where appropriate. HPLC purification was carried out using a Shimadzu LC-20 AD system, and elution profiles monitored using a Shimadzu SPD-10A.

#### *Abbreviations*

Ac, acetyl; Cys, cysteine; Et, ethyl; DIAD, diisopropyl azodicarboxylate; DIBAL-H, diisobutylaluminium hydride; DMAP, 4-dimethylaminopyridine; DMF, *N,N*-dimethylformamide; DMSO, dimethyl sulfoxide; HMPA, hexamethylphosphoric triamide; Me, methyl Ms, mesyl; NaHMDS, sodium hexamethylenedisilane; MOM, methoxymethyl; MTPA, α-methoxy-α-(trifluoromethyl)phenylacetyl; NMP, *N*-methylpyrrolidone; rt, room temperature; TBAF, tetrabutylammonium fluoride; TBDPS, *tert*-butyldiphenylsilyl; TBS, *tert*-butyldimethylsilyl; THF, tetrahydrofuran; TMS, trimethylsilyl; Ts, tosyl.

***Outline of the synthesis of S11***

The synthetic route to **S11** is outlined in Scheme S1. **S7** was synthesized according to the procedures described in *ref S1*. **S8** was synthesized from 1-pentyne by tributyl stannylation using tributyltin hydride from 1-pentyne. **S9** was synthesized by a Stille coupling reaction using **S7** and **S8**. Removing TBDPS from **S9** provided **S10**, which was subsequently subjected to Parikh–Doering oxidation to give aldehyde **S11**.

**Scheme S1.**

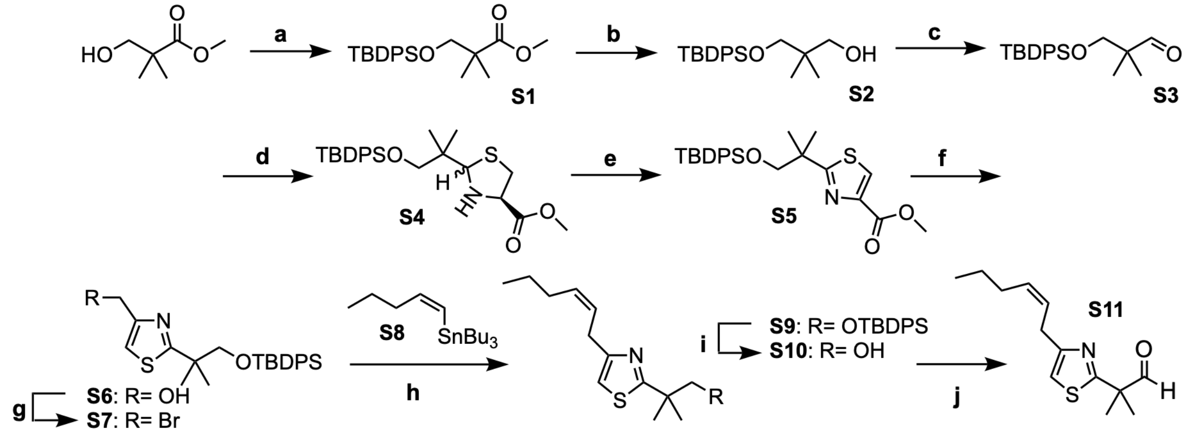


*Reagents and conditions*; (a) TBDPSCl, imidazole, DMF, 0 ˚C to rt, 14 h; (b) DIBAL-H, $Et_2O$, -78 ˚C, 30 min, 92% (over 2 steps); (c) pyridine·$SO_3$, DMSO, $Et_3N$, $CH_2Cl_2$, 0 ˚C to rt, 30 min, quant.; (d) L-Cys-OMe·HCl, $Et_3N$, toluene, 0 ˚C to rt, 24 h, 62%; (e) $MnO_2$, pyridine, benzene, reflux, 12 h, 58%; f) DIBAL-H, $Et_2O$, -78 ˚C, 30 min, 70%; (g) 1) $Ms_2O$, $Et_3N$, $CH_2Cl_2$, 0 ˚C, 20 min, 2) acetone, LiBr, rt, 1 h, 80%; (h) **S8**, $Pd(MeCN)_2Cl_2$(10 mol %), NMP, rt, 10 min, 77%; (i) TBAF·$3H_2O$, THF, 55 ˚C, 22 h, quant.; (j) pyridine·$SO_3$, $Et_3N$, DMSO, $CH_2Cl_2$, 0 ˚C to rt, 2 h, 87%.

**Synthesis of S1:** To an ice-cooled solution of methyl hydroxypivalate (5.0 g, 37.8 mmol) in DMF (75 mL), TBDPSCl (9.96 g, 38.9 mmol) and imidazole (6.18 g, 90.7 mmol) were added, and the mixture was stirred at room temperature for 3 h. The reaction mixture was quenched with saturated aqueous $NH_4Cl$, extracted with $Et_2O$, washed with water and brine, and dried over anhydrous $MgSO_4$. The resulting mixture was filtered and concentrated *in vacuo* to afford crude **S1** as a colorless oil (13.3 g, containing biproducts derived from TBDPS-Cl), which was used for the next step without further purification: $^1$H-NMR (400 MHz, $CDCl_3$) $\delta$ 7.64 (m, 4H), 7.40 (m, 6H), 3.67 (s, 3H), 3.64 (s, 2H), 1.20 (s, 6H), 1.03 (s, 9H).

**Synthesis of S2:** To solution of **S1** (13.3 g) in $Et_2O$ (70 mL), DIBAL-H (in *n*-hexane 87.2 mL, 89.8 mmol) was added dropwise at -78 ˚C, and the mixture was stirred at -78 ˚C for 20 min under a nitrogen atmosphere. The reaction mixture was diluted with $Et_2O$ (70 mL) and quenched with saturated aqueous Rochelle salt (potassium sodium tartrate) solution and then stirred for 1 h at room temperature. The mixture was extracted with ethyl acetate, washed with water and brine, and dried over anhydrous $Na_2SO_4$. The extract was concentrated *in vacuo* to afford crude **S2** as a colorless oil (11.9 g, 34.8 mmol, 92%, over 2steps), which was used for the next step without further purification: $^1$H-NMR (400 MHz, $CDCl_3$) $\delta$ 7.66 (m, 4H), 7.40 (m, 6H), 3.51 (d, $J$ = 5.2 Hz, 2H), 3.48 (s, 2H), 2.41 (t, $J$ = 5.2 Hz, 1H), 1.07 (s, 9H), 0.89 (s, 6H).

**Synthesis of S3:** To an ice-cooled solution of **S2** (5.5 g, 16.1 mmol) in $CH_2Cl_2$ (45 mL), $Et_3N$ (6.4 mL, 46.4 mmol) and pyridine·$SO_3$ (7.4 g, 46.4 mmol) in DMSO were added under a nitrogen atmosphere. After stirring mixture at room temperature for 30 min, it was quenched with saturated aqueous $NaHCO_3$, and $CH_2Cl_2$ was removed *in vacuo*. The resulting mixture was extracted with $Et_2O$, washed with $NH_4Cl$ and brine, and the organic layer was dried over anhydrous $MgSO_4$. The resulting extract was concentrated *in vacuo* to afford crude **S3** as a pale-yellow oil (5.5 g, 16.1 mmol, quant.), which was used for the next step without further purification: $^1$H-NMR (400 MHz, $CDCl_3$) $\delta$ 9.60 (s, 1H), 7.67 (m, 4H), 7.41 (m, 6H), 3.64 (s, 2H), 1.07 (s, 6H), 1.03 (s, 9H).

**Synthesis of S4:** To an ice-cooled solution of **S3** (5.5 g, 16.1 mmol) in toluene, (*S*)-H-Cys-OMe·HCl (3.02 g, 17.6 mmol) and $Et_3N$ (2.55 mL, 18.4 mmol) were added under a nitrogen atmosphere. After the reaction mixture was stirred at room temperature for 22 h, it was concentrated *in vacuo*. The residue was purified by silica-gel column chromatography (Wako gel® C-200, 10% Ethyl acetate/*n*-hexane) to afford **S4** as a colorless oil (4.86 g, 10.6 mmol, 62%): ESI-MS (*m/z*) 458.2 $[M+H]^+$, 480.2 $[M+Na]^+$. As **S4** was a mixture of diastereomers to give extremely complicated NMR spectra, we did not annotate the spectra.

**Synthesis of S5:** A suspension $MnO_2$ (1.0 g) in benzene (25 mL) was heated under reflux for 3 h on a Dean-Stark apparatus to remove water. Then, **S4** (575 mg, 1.25 mmol) in benzene (5 mL) and pyridine (255 μL) were added to the suspension, followed by the stirring for 12 h under reflux. The reaction mixture was cooled to room temperature and filtered with a pad of celite. The filtrate was concentrated *in vacuo*, and the residue was purified by silica-gel column chromatography (Wako gel® C-200, 10–20% Ethyl acetate/*n*-hexane) afforded **S5** as a colorless oil (328 mg, 0.723 mmol, 58%): $^1$H-NMR (400 MHz, $CDCl_3$) $\delta$ 8.09 (s, 1H), 7.56 (m, 4H), 7.38 (m, 6H), 3.94 (s, 3H), 3.75 (s, 2H), 1.48 (s, 6H), 1.00 (s, 9H); $^{13}$C-NMR (100 MHz, $CDCl_3$) $\delta$ 178.96, 162.45, 146.29, 135.84 (4C), 133.39 (2C), 129.88 (2C), 127.86 (4C), 127.42, 72.58, 52.52, 43.63, 27.00 (3C), 25.61 (2C), 19.51.

**Synthesis of S6:** To a solution of **S5** (1.6 g, 3.53 mmol) in $Et_2O$ (20 mL) at -78 ˚C, DIBAL-H (in *n*-hexane 10 mL, 10.3 mmol) was added dropwise under a nitrogen atmosphere, and the mixture was stirred at -78 ˚C for 30 min. The reaction mixture was diluted with $Et_2O$ (20 mL) and quenched with saturated aqueous Rochelle salt solution (40 mL), and stirred 1 h at room temperature. The mixture was extracted with $Et_2O$, washed with water and brine, and dried over anhydrous $MgSO_4$. The purification of the crude product by silica-gel column chromatography (Wako gel® C-200, 40% ethyl acetate/*n*-hexane) afforded **S6** as a colorless solid (1.05 g, 2.47 mmol, 70%): $^1$H-NMR (400 MHz, $CDCl_3$) $\delta$ 7.56 (m, 4H), 7.38 (m, 6H), 7.04 (s, 1H), 4.73 (s, 2H), 3.73 (s, 2H), 2.20 (brs, 1H), 1.45 (s, 6H), 1.00 (s, 9H).

**Synthesis of S7:** To an ice-cooled solution of **S6** (300 mg, 0.705 mmol) in $CH_2Cl_2$ (2.3 mL), $Et_3N$ (148 μL, 1.06 mmol) and $Ms_2O$ (160 mg, 0.92 mmol) were added under a nitrogen atmosphere, followed by the stirring the mixture at 0 ˚C for 20 min. Then, acetone (3.0 mL) and LiBr (367 mg, 4.23 mmol) were added to the mixture, and the mixture was further stirred at room temperature for 1 h to convert the intermediate to the desired bromide. The reaction mixture was quenched with saturated aqueous $NH_4Cl$, extracted with $CH_2Cl_2$, and the aqueous phase was further extracted with $Et_2O$. The combined organic phases were washed with water and brine, and dried over anhydrous $MgSO_4$. The crude product was purified by silica-gel column chromatography (Wako gel® C-200, 10% ethyl acetate/*n*-hexane) to afford **S7** as a colorless oil (275 mg, 0.563 mmol, 80%): $^1$H-NMR (400 MHz, $CDCl_3$) $\delta$ 7.58 (m, 4H), 7.38 (m, 6H), 7.19 (s, 1H), 4.57 (s, 2H), 3.73 (s, 2H), 1.44 (s, 6H), 0.99 (s, 9H); $^{13}$C-NMR (100 MHz, $CDCl_3$) $\delta$ 178.83, 151.54, 135.86 (4C), 133.57 (2C), 129.81 (2C), 127.84 (4C), 117.15, 72.81, 43.44, 27.84, 26.98 (3C), 25.56 (2C), 19.53.

**Synthesis of S8:** To an ice-cooled solution of zirconium (IV) chloride (0.23 g, 1.0 mmol) in *n*-hexane (2.5 mL), 1-pentyne (0.49 mL, 5.0 mmol) was added under a nitrogen atmosphere. The mixture was stirred at 0 ˚C for 5 h, followed by the addition of tributyltin hydride (2.0 mL, 7.5 mmol). Stirring was continued for 1 h at 0 °C, and $Et_3N$ (0.35 mL, 2.5 mmol) was then added. The reaction mixture was stirred at room temperature for an additional 5 min and filtered through a pad of Celite. The filtrate was concentrated *in vacuo*, and the residue was purified by silica-gel column chromatography (Wako gel® C-200, *n*-hexane) to afford **S8** as a colorless oil containing inseparable biproducts (2.19 g): $^1$H-NMR (400 MHz, $CDCl_3$) $\delta$ 6.51 (dt, $J$ = 7.0, 12.4 Hz, 1H), 5.79 (dt, $J$ = 12.4, 1.1 Hz, 1H), 2.00 (ddt, $J$ = 7.4, 7.2, 1.1 Hz, 2H), 1.56-1.25 (m, 14H), 0.94-0.87 (m, 18H); $^{13}$C-NMR (100 MHz, $CDCl_3$) $\delta$ 149.31, 128.13, 39.38, 29.44 (3C), 27.56 (3C), 23.26, 13.92, 10.47, 8.99.

**Synthesis of S9:** To a solution of **S7** (111 mg, 0.310 mmol) and **S8** (138 mg, 0.282 mmol) in nitrogen-purged NMP (1.4 mL), $Pd(MeCN)_2Cl_2$ (7.32 mg, 0.0282 mmol) was added under a nitrogen atmosphere. After stirring at room temperature for 10 min, the reaction mixture was quenched with aqueous ammonia (28% (w/v), 1.65 mL) and at room temperatur stirred for additional 30 mine. The mixture was partitioned with ethyl acetate and saturated aqueous Rochelle salt solution. The organic phase was washed with brine and dried over anhydrous $Na_2SO_4$. The crude product was purified by silica gel column chromatography (Wako Gel® C-200, 10% ethyl acetate/*n*-hexane) to afford **S9** as a pale yellow oil (104 mg, 0.218 mmol, 77% over two steps): $^1$H-NMR (500 MHz, $CDCl_3$) $\delta$ 7.57 (m, 4H), 7.38 (m, 6H), 6.75 (s, 1H), 5.72-5.54 (m, 2H), 3.73 (s, 2H), 3.55 (d, $J$ = 7.2 Hz, 2H), 2.10 (dt, $J$ = 7.2, 7.2 Hz, 2H), 1.30-1.45 (m, 2H), 1.45 (s, 6H), 0.99 (s, 9H), 0.91 (t, 3H); $^{13}$C-NMR (125 MHz, $CDCl_3$) $\delta$ 177.68, 155.58, 135.87 (4C), 133.74 (2C), 131.95, 129.75 (2C), 127.78 (4C), 126.25, 112.00, 72.97, 43.19, 29.99, 29.55, 26.95 (3C), 25.61 (2C), 22.97, 19.54, 13.82.

**Synthesis of S10:** To a solution of **S9** (930 mg, 1.95 mmol) in THF (9 mL), TBAF·$3H_2O$ (1.30 g, 4.10 mmol) was added under a nitrogen atmosphere, and the mixture was stirred at 55 ˚C for 22 h. The reaction mixture was quenched with water, extracted with ethyl acetate. The organic phase was washed with water and brine and dried over anhydrous $Na_2SO_4$. The crude product was purified by silica-gel column chromatography (Wako gel® C-200, 0–20% Ethyl acetate/*n*-hexane) to afford **S10** as a colorless oil (460 mg, 1.92 mmol, quant.): $^1$H-NMR (400 MHz, $CDCl_3$) $\delta$ 6.76 (s, 1H), 5.70-5.50 (m, 2H), 4.30 (brs, 1H), 3.69 (s, 2H), 3.51 (d, $J$ = 6.8 Hz, 2H), 2.10 (dt, $J$ = 7.1, 7.1 Hz, 2H), 1.45-1.30 (m, 2H), 1.39 (s, 6H), 0.91 (t, $J$ = 7.4 Hz, 3H); $^{13}$C-NMR (125 MHz, $CDCl_3$) $\delta$ 179.2, 155.42, 132.18, 125.84, 111.75, 71.84, 42.17, 29.52, 29.52, 26.60 (2C), 22.93, 13.99.

**Synthesis of S11:** To an ice-cooled solution of **S10** (72 mg, 0.301 mmol) in $CH_2Cl_2$ (1.2 mL), $Et_3N$ (168 μL, 1.20 mmol) and pyridine·$SO_3$ (192 mg, 1.20 mmol, dissolved in 1.2 mL DMSO) were added under a nitrogen atmosphere. The mixture was stirred at room temperature for 1.5 h and then quenched with saturated aqueous $NaHCO_3$. $CH_2Cl_2$ was removed *in vacu*o, and the residue was extracted with $Et_2O$. The organic phase was washed with saturated aqueous $NH_4Cl$ and brine and dried over anhydrous $MgSO_4$. The crude product was purified by silica-gel column chromatography (Wako gel® C-200, 10% Ethyl acetate/*n*-hexane) to afford **S11** as a colorless oil (62 mg, 0.261 mmol, 87%): $^1$H-NMR (500 MHz, $CDCl_3$) $\delta$ 9.69 (s, 1H), 6.85 (s, 1H), 5.70-5.50 (m, 2H), 3.56 (d, $J$ = 10 Hz, 2H), 2.10 (dt, $J$ = 7.2 Hz, 2H), 1.57 (s, 6H), 1.42 (tq, $J$ = 7.2, 7.2 Hz, 2H), 0.92 (t, $J$ = 7.5 Hz, 3H); $^{13}$C-NMR (125 MHz, $CDCl_3$) $\delta$ 200.31, 171.68, 157.05, 132.31, 125.76, 113.26, 52.00, 29.86, 29.53, 23.11 (2C), 22.91, 13.99.

#### *Outline of the synthesis of MT1*

The synthetic route to **MT1** is outlined in Scheme S2. **S16** and the chiral auxiliary **S17** were synthesized according to the procedures described in *refs S1 and S2*, respectively. An Evans aldol reaction of **S11** with the chiral auxiliary **S17** (*ref. S3*) gave **S18**. Protection of hydroxy group of **S18** as a TBS ether gave **S19**, which was subsequently reduced with DIBAL-H provided **S20**. **S21** was synthesized from **S20** via Horner–Wadsworth–Emmons reaction with $Ph_3P^+CH_2I{\cdot}I^-$. **MT1** was synthesized by a Stille coupling reaction between **S16** and **S21**, followed by deprotection of TBS group. To confirm the stereochemistry of **MT1**, its absolute configuration was determined using the Mosher ester method. Derivatization of **MT1** with (*R*)- and (*S*)-MTPA-Cl afforded **S23** and **S24**, respectively. Comparison of the $^1$H-NMR spectra of these compounds and analysis of the corresponding chemical-shift differences confirmed that **MT1** possesses the intended *R* configuration.

**Scheme S2.**

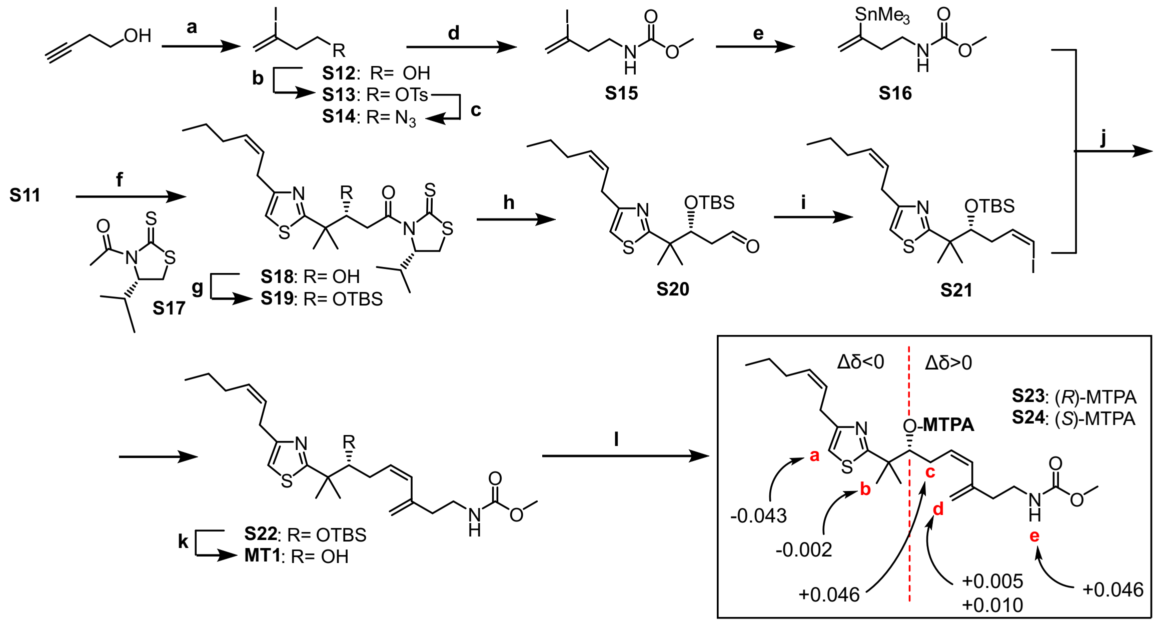


*Reagents and conditions*; (a) TMSCl, NaI, $H_2O$, $CH_3CN$, rt, 1 h; (b) TsCl, $Et_3N$, $Me_3N{\cdot}HCl$, $CH_2Cl_2$, 0 ˚C, 2 h, 45% (2 steps); (c) $NaN_3$, DMF, 0 ˚C, 2 h, then 50 ˚C 1.5 h; (d) 1) $Ph_3P$, $H_2O$, THF, rt, 16 h, 2) metyl chloroformate, $Et_3N$, rt, 4 h, 62% (2 steps); (e) hexamethylditin, $Pd(CH_3CN)_2Cl_2$, NMP, rt, 4 h, 58%; (f) **S17**, $TiCl_4$, *i*$Pr_2NEt$, $CH_2Cl_2$, -50 ˚C, 2 h, 82%; (g) TBSOTf, 2,6-lutidine, $CH_2Cl_2$, 0 ˚C to rt, 2 h, 91%; (h) DIBAL-H, toluene, -78 ℃, 1 h, 69%; (i) $Ph_3P^+CH_2I{\cdot}I^-$, NaHMDS, HMPA, THF, -78 ℃, 20 min, 96%; (j) CuI, $Pd(Ph_3P)_4$, CsF, DMF, rt, 40 min, 89%; (k) TBAF·$3H_2O$, THF, 0 ˚C to rt, 2 h, 42%; (l) (*R*)-MTPA-Cl, DMAP, pyridine, THF, rt, 26 h, quant. (for **S23**); (*S*)-MTPA-Cl, DMAP, pyridine, THF, rt, 27 h, quant.(for **S24**).

**Synthesis of S13:** To a solution of NaI (6.00 g, 40.0 mmol) and TMSCl (5.1 mL, 40.0 mmol) in $CH_3CN$ (30 mL), water (0.36 mL, 20.0 mmol) was added. The mixture was stirred at room temperature for 10 min, followed by the addition of 3-butyn-1-ol (1.5 mL, 20.0 mmol) in $CH_3CN$ (5.0 mL). Stirring was continued for 1 h at room temperature, and the reaction was then quenched with water. The mixture was extracted with $Et_2O$, and the organic phase was dried over anhydrous $MgSO_4$. Removal of the solvent *in vacuo* afforded iodide **S12**, which was used directly in the next reaction without further purification.

To an ice-cooled solution of **S12** in $CH_2Cl_2$ (25 mL), TsCl (5.30 g, 27.8 mmol), $Et_3N$ (5.58 mL, 40.0 mmol), and $Me_3N \cdot H_2O$ (0.19 g, 2.0 mmol) were added. The mixture was stirred at 0 °C for 2 h and then quenched with water. The mixture was extracted with $Et_2O$, and the organic phase was dried over anhydrous $MgSO_4$. The crude product was purified by silica gel column chromatography (Wako Gel® C-200, 15–40% ethyl acetate/*n*-hexane). The fractions containing **S13** were combined and washed successively with 1.0 M aqueous $KHSO_4$, saturated aqueous $Na_2S_2O_3$, water, and brine, and then dried over anhydrous $MgSO_4$. Removal of the solvent afforded **S13** as a brown oil (3.17 g, 9.00 mmol, 45% over 2 steps): $^1$H-NMR (400 MHz, $CDCl_3$) $\delta$ 7.85-7.78 (m, 2H), 7.35 (d, $J$ = 8.0 Hz, 2H), 6.11 (d, $J$ = 1.6 Hz, 1H), 5.78 (d, $J$ = 1.8 Hz, 1H), 4.14-4.08 (t, $J$ = 6.2 Hz, 2H), 2.71 (t, $J$ = 6.0 Hz, 2H), 2.45 (s, 3H); $^{13}$C-NMR (100 MHz, $CDCl_3$) $\delta$ 145.15, 133.07, 130.11 (2C), 129.42, 128.24 (2C), 103.79, 68.29, 44.59, 21.87; ESI-MS (*m/z*) 374.9 $[M+Na]^+$.

**Synthesis of S15:** To an ice-cooled solution of **S13** (3.17 g, 9.00 mmol) in DMF (23.4 mL), sodium azide (1.76 g, 27.0 mmol) was added under a nitrogen atmosphere, and the mixture was stirred at 0 °C for 2 h. The reaction mixture was then heated to 50 °C and stirred for an additional 3 h. The reaction was quenched with water, and the mixture was extracted with $Et_2O$. The organic phase was washed successively with 1 M aqueous $KHSO_4$, water, and brine and dried over anhydrous $MgSO_4$. Removal of the solvent *in vacuo* afforded crude **S14**, which was used directly in the next reaction without further purification.

To an ice-cooled solution of azide **S14** in THF (29 mL), triphenylphosphine (3.45 g, 13.2 mmol) was added, and the mixture was stirred at room temperature for 2 h. Water (0.24 mL, 13 mmol) was subsequently added, and stirring was continued for an additional 18 h at room temperature to afford the corresponding amine. The reaction mixture was cooled to 0 °C, followed by the addition of methyl chloroformate (1.35 mL, 17.6 mmol) and $Et_3N$ (2.1 mL, 15.1 mmol), and then stirred at room temperature for 4 h. The reaction was quenched with water, and the mixture was extracted with $Et_2O$. The organic phase was washed with 1 M aqueous $KHSO_4$ and water and dried over anhydrous $MgSO_4$. The crude product was purified by silica gel column chromatography (Wako Gel® C-200, 10–40% ethyl acetate/*n*-hexane) to afford **S15** as a colorless oil (1.43 g, 5.60 mmol, 62% over two steps): $^1$H-NMR (400 MHz,

$CDCl_3$) $\delta$ 6.11 (br t, $J$ = 1.3 Hz, 1H), 5.81 (br d, $J$ = 1.4 Hz, 1H), 4.77 (br s, 1H), 3.66 (s, 3H), 3.37-3.32 (m, 2H), 2.58 (t, $J$ = 5.6 Hz, 2H); $^{13}$C-NMR (100 MHz, $CDCl_3$) $\delta$ 157.14, 128.38, 108.00, 52.36, 45.36, 40.100; ESI-MS (*m/z*) 256.0 $[M+H]^+$, 293.9 $[M+K]^+$.

**Synthesis of S16:** To an ice-cooled solution of **S15** (0.20 g, 0.80 mmol) in argon-purged NMP (5 mL), hexamethylditin (0.52 g, 1.6 mmol) and $Pd(MeCN)_2Cl_2$ (21 mg, 0.08 mmol) were added sequentially. The mixture was stirred for 2 h at room temperature and then for an additional at 30 °C 2 h to complete the reaction. The reaction was quenched with water, and the mixture was extracted with $Et_2O$. The organic phase was washed successively with 1.0 M aqueous $KHSO_4$, water, and brine and dried over anhydrous $MgSO_4$. The crude product was purified by silica gel column chromatography (Wako Gel® C-200, 15–40% ethyl acetate/*n*-hexane) to afford **S16** as a pale brown oil (75 mg, 0.26 mmol, 32%): $^1$H-NMR $^1$H-NMR (400 MHz, $CDCl_3$) $\delta$ 5.72 (dt, $J$ = 2.5, 1.4 Hz, 1H), 5.27 (dt, $J$ = 2.6, 0.9 Hz, 1H), 4.63 (br s, 1H), 3.66 (s, 3H), 3.26-3.22 (m, 2H), 2.46 (t, $J$ = 6.7 Hz, 2H), 0.16 (s, 9H); $^{13}$C-NMR (100 MHz, $CDCl_3$) $\delta$ 156.89, 152.25, 127.41, 52.03, 40.59, 40.19, -9.50 (3C); ESI-MS (*m/z*) 294.1 $[M+H]^+$, 316.0 $[M+Na]^+$.

**Synthesis of S17:** This compound was synthesized according to the procedure described in *ref. S3*.

**Synthesis of S18:** To a solution of **S17** (269 mg, 1.32 mmol) in $CH_2Cl_2$ (6.4 mL), $TiCl_4$ (155 μL, 1.41 mmol) was added at -50 °C under a nitrogen atmosphere. The mixture was stirred at -50 °C for 20 min, followed by the addition of *i*-$Pr_2NEt$ (240 μL, 1.41 mmol), and stirring was continued at -40 °C for 1 h. The reaction mixture was then cooled to −78 °C, and a solution of **S11** (209 mg, 0.881 mmol) in $CH_2Cl_2$ (1.2 mL) was added. The mixture was stirred at -40 °C for 100 min and then quenched with phosphate buffer and Rochelle salt solution. The mixture was extracted with $CH_2Cl_2$, and the organic phase was washed with brine and dried over anhydrous $MgSO_4$. The crude product was purified by silica gel column chromatography (Wako Gel® C-200, 10–40% ethyl acetate/*n*-hexane) to afford **S18** as a yellow oil (320 mg, 0.726 mmol, 82%): $^1$H-NMR (500 MHz, $CDCl_3$) $\delta$ 6.77 (s, 1H), 5.70-5.50 (m, 2H), 5.14 (dt, $J$ = 0.8 Hz, 7.0 Hz, 1H), 4.77 (brs, 1H), 4.34 (brt, 1H), 3.45~3.56 (m, 3H), 3.38 (d, $J$ = 6.3 Hz, 2H), 3.01 (dd, $J$ = 1.0 Hz, 11.4 Hz, 1H), 2.45-2.30 (m, 1H), 2.10 (dt, $J$ =7.1, 7.1 Hz, 2H), 1.47 (s, 3H), 1.45-1.35 (m, 2H), 1.43 (s, 3H), 1.06 (d, $J$ = 6.8 Hz, 3H), 0.98 (d, $J$ = 6.9 Hz, 3H), 0.92 (t, $J$ = 7.4 Hz, 3H); $^{13}$C-NMR (125 MHz, $CDCl_3$) $\delta$ 203.38, 178.10, 172.86, 155.86, 132.15, 125.89, 112.00, 75.39, 72.10, 44.12, 41.00, 30.55, 29.76, 29.53, 26.86, 26.4, 24.72, 22.93, 19.32, 17.92, 14.00; ESI-MS (*m/z*) 441.1 $[M+H]^+$.

**Synthesis of S19:** To an ice-cooled solution of **S18** (300 mg, 0.681 mmol) in $CH_2Cl_2$ (6.4 mL), 2,6-lutidine (143 μL, 1.23 mmol) and TBSOTf (251 μL, 1.09 mmol) were added under a nitrogen atmosphere, and the mixture was stirred at 0 °C for 30 min. The reaction was quenched with saturated aqueous $NH_4Cl$, and the mixture was extracted with $Et_2O$. The organic phase was washed successively with Rochelle salt solution, water, and brine and dried over anhydrous $MgSO_4$. The crude product was purified by silica gel column chromatography (Wako Gel® C-200, 20% ethyl acetate/*n*-hexane) to afford **S19**as a yellow oil (343 mg, 0.618 mmol, 91%): $^1$H-NMR (400 MHz, $CDCl_3$) $\delta$ 6.72 (s, 1H), 5.71-5.51 (m, 2H), 5.08 (dt, *J* = 0.8, 7.0 Hz, 1H), 4.74 (t, *J* = 4.8 Hz, 1H), 3.56-3.45 (m, 3H), 3.35 (d, *J* = 6.3 Hz, 2H), 3.00 (dd, *J* = 1.0 Hz, 11.4 Hz, 1H), 2.40-2.25 (m, 1H), 2.09 (dt, *J* = 7.1, 7.1 Hz, 2H), 1.45-1.35 (m, 2H), 1.44 (s, 3H), 1.37 (s, 3H), 1.04 (d, *J* = 6.8 Hz, 3H), 0.94 (d, *J* = 6.9 Hz, 3H), 0.91 (t, *J* = 7.4 Hz, 3H), 0.85 (s, 9H), 0.03 (s, 3H), -0.01 (s, 3H); $^{13}$C-NMR (125 MHz, $CDCl_3$) $\delta$ 555.2, 202.85, 177.28, 172.32, 155.55, 131.85, 126.30, 112.25, 75.51, 71.75, 46.13, 43.62, 31.12, 30.92, 29.97, 29.55, 26.18 (3C), 25.86, 25.47, 22.97, 19.31, 18.48, 18.22 (2C), 14.03, -2.70, -4.30; ESI-MS (*m/z)* 555.2 $[M+H]^+$.

**Synthesis of S20:** To a solution of **S19** (5.0 g, 37.8 mmol) in toluene (6 mL) at -78 ˚C under Ar atmosphere, DIBAL-H (in n-hexane 1.50 mL, 1.54 mmol) was added slowly, and the mixture was stirred for at -78 °C 1 h. The reaction mixture was quenched with Rochelle salt solution, and the mixture was stirred for 1 h at room temperature. The mixture was extracted with $Et_2O$, washed with Rochelle salt solution and brine, and dried over anhydrous $MgSO_4$. After filtration and concentration *in vacuo*, the purification by silica-gel column chromatography (Wako gel® C-200, 10–20% Ethyl acetate/*n*-hexane) afforded **S20** as a pale yellow oil (168 mg, 0.424 mmol, 69%): $^1$H-NMR (400 MHz, $CDCl_3$) $\delta$ 9.58 (s, 1H), 6.75 (s, 1H), 5.70-5.50 (m, 2H), 4.56 (t, *J* = 5.2 Hz, 1H), 3.52 (d, *J* = 7.2 H, 2H), 2.70-2.40 (m, 2H), 2.09 (dt, *J* = 7.1, 7.1 Hz, 2H), 1.45-1.35 (m, 2H), 1.43 (s, 3H), 1.38 (s, 3H), 0.92 (t, *J* = 7.4 Hz, 3H), 0.86 (s, 9H), 0.03 (s, 3H), -0.01 (s, 3H); $^{13}$C-NMR (100 MHz, $CDCl_3$) $\delta$ 201.16, 176.82, 155.78, 132.02, 126.10, 112.57, 74.19, 48.77, 45.99, 29.92, 29.54, 26.08 (3C), 25.92, 24.41, 22.95, 18.32, 14.02, -2.73, -4.66; ESI-MS (*m/z*) 496.2 $[M+H]^+$.

**Synthesis of S21:** To a solution of $Ph_3P^+CH_2I \cdot I^-$ (1.12 g, 2.12 mmol) in THF (6 mL), NaHMDS (2.12 mL, 2.12 mmol) was added under an argon atmosphere, and the mixture was stirred at room temperature for 5 min. The reaction mixture was then cooled to −78 °C, and HMPA (738 μL, 4.24 mmol) was added. After stirring at -78 °C for 5 min, **S20** (168 mg, 0.424 mmol) was added, and stirring was continued for an additional 20 min at -78 °C. The reaction was quenched with water at −78 °C and then allowed to warm to room temperature. The mixture was extracted with $Et_2O$, and the organic phase was washed with water and brine and dried over anhydrous $MgSO_4$. The crude product was purified by silica gel column chromatography (Wako Gel® C-200, 0–10% ethyl acetate/*n*-hexane) to afford **S21** as a colorless oil (212 mg, 0.408 mmol, 96%): $^1$H-NMR (400 MHz, $CDCl_3$) $\delta$ 6.76 (s, 1H), 6.17-6.05 (m, 2H), 5.70-5.50 (m, 2H), 4.19 (t, $J$ = 5.3 Hz, 1H), 3.53 (d, $J$ = 7.2 Hz, 2H), 2.37-2.23 (m, 2H), 2.09 (dt, $J$ = 7.1, 7.1 Hz, 2H), 1.45-1.35 (m, 2H), 1.43 (s, 3H), 1.38 (s, 3H), 0.91 (t, $J$ = 7.4 Hz, 3H), 0.86 (s, 9H), 0.03 (s, 3H), -0.01 (s, 3H); $^{13}$C-NMR (100 MHz, $CDCl_3$) $\delta$ 177.98, 155.63, 138.98, 131.94, 126.25, 112.10, 82.86, 78.12, 46.30, 39.60, 30.00, 29.56, 26.20 (3C), 25.92, 24.69, 22.97, 18.39, 14.04, -2.73, -4.66; ESI-MS (*m/z*) 520.1 $[M+H]^+$.

**Synthesis of S22:** To an argon-purged solution of **S21** (42 mg, 0.080 mmol) and **S16** (42 mg, 0.14 mmol) in DMF (0.5 mL), CuI (3 mg, 0.016 mmol), $Pd(PPh_3)_4$ (5 mg, 0.004 mmol), and CsF (24 mg, 0.16 mmol) were added successively. The mixture was stirred at room temperature for 10 min, quenched with water, and extracted with $Et_2O$. The organic phase was washed with water and dried over anhydrous $MgSO_4$. The crude product was purified by silica gel column chromatography (Wako Gel® C-200, 20% ethyl acetate/*n*-hexane) to afford **S22** as a yellow oil (37 mg, 0.071 mmol, 89%): $^1$H-NMR (400 MHz, $CDCl_3$) $\delta$ 6.72 (s, 1H), 5.72 (m, 1H), 5.70-5.50 (m, 4H), 5.26 (m, 1H), 4.63 (br s, 1H), 4.05 (t, $J$ = 5.2 Hz, 1H), 3.65 (s, 3H), 3.51 (d, $J$ = 6.8 Hz, 2H), 3.23 (m, 2H), 2.46 (t, $J$ = 6.7 Hz, 2H), 2.43-2.28 (m, 2H), 2.08 (dt, $J$ = 7.1, 7.1 Hz, 2H), 1.45-1.35 (m, 2H), 1.40 (s, 3H), 1.34 (s, 3H), 0.91 (t, $J$ = 7.4 Hz, 3H), 0.86 (s, 9H), 0.03 (s, 3H), -0.01 (s, 3H); $^{13}$C-NMR (100 MHz, $CDCl_3$) $\delta$ 178.24, 157.14, 155.51, 142.12, 131.9, 131.00, 129.24, 126.24, 116.00, 112.08, 79.62, 52.22, 46.44, 39.55, 37.55, 33.31, 29.91, 29.52, 26.52 (3C), 26.06, 24.75, 22.94, 18.44, 14.00, -3.38, -4.46; ESI-MS (*m/z*) 521.3 $[M+H]^+$.

**Synthesis of MT1:** To an ice-cooled solution of **S22** (19 mg, 0.036 mmol) in THF (1.0 mL), $TBAF\cdot 3H_2O$ (61 mg, 0.194 mmol) was added, and the mixture was stirred at room temperature for 2 h. The reaction was quenched with water, and the mixture was extracted with ethyl acetate. The organic phase was washed with brine and dried over anhydrous $Na_2SO_4$. The crude product was purified by silica gel column chromatography (Wako Gel® C-200, 20–40% ethyl acetate/*n*-hexane) to afford **MT1** as a pale yellow oil (6 mg, 0.015 mmol, 42%): $^1$H-NMR (400 MHz, $CDCl_3$) $\delta$ 6.76 (s, 1H), 5.87 (d, $J$ = 11.8 Hz, 1H), 5.70-5.50 (m, 3H), 5.44 (brs, 1H), 5.01 (s, 1H), 4.94 (brs, 1H), 4.88 (s, 1H), 4.63 (brs, 1H), 3.78 (d, $J$ = 9.8 Hz, 1H), 3.62 (s, 3H), 3.50 (d, $J$ = 6.8 Hz, 2H), 3.40-3.10 (m, 2H), 2.50-2.15 (m, 4H), 2.09 (dt, $J$ = 7.1, 7.1 Hz, 2H), 1.45-1.34 (m, 2H), 1.43 (s, 3H), 1.38 (s, 3H), 0.91 (t, $J$ = 7.4 Hz, 3H); $^{13}$C-NMR (100 MHz, $CDCl_3$) $\delta$ 179.53, 157.36, 155.65, 142.73, 132.22, 131.09, 130.83, 125.79, 116.05, 111.82, 78.36, 52.04, 44.77, 39.61, 37.38, 31.80, 29.72, 29.5, 26.89, 24.03, 22.92, 13.99; ESI-MS (*m/z*) 407.2 $[M+H]^+$.

***Determination of the absolute configuration of MT1 by Mosher ester method***

**Synthesis of S23:** To a solution of **MT1** (2.5 mg, 0.0061 mmol) in $CH_2Cl_2$ (0.3 mL), pyridine (1.47 µL, 0.0183 mmol), DMAP (2.24 mg, 0.0183 mmol), and (*S*)-MTPA-Cl (2.28 µL, 0.0122 mmol) were added successively, and the mixture was stirred at room temperature for 3 h. To ensure complete derivatization, additional pyridine (12 µL, 0.146 mmol), DMAP (18.4 mg, 0.146 mmol), and (*S*)-MTPA-Cl (18.4 µL, 0.0976 mmol) were added, and stirring was continued at room temperature for an additional 27 h. The reaction was quenched with water, and the mixture was extracted with ethyl acetate. The organic phase was washed with brine and dried over anhydrous $MgSO_4$. The crude product was purified by silica gel column chromatography (Wako Gel® C-200, 50% ethyl acetate/*n*-hexane) to afford **S23** as a colorless oil (4.0 mg, quant.).

**Synthesis of S24:** To a solution of **MT1** (2.5 mg, 0.0061 mmol) in $CH_2Cl_2$ (0.3 mL), pyridine (1.47 µL, 0.0183 mmol), DMAP (2.24 mg, 0.0183 mmol), and (*R*)-MTPA-Cl (2.28 µL, 0.0122 mmol) were added successively, and the mixture was stirred at room temperature for 3 h. To ensure complete derivatization, additional pyridine (5.9 µL, 0.0732 mmol), DMAP (9.2 mg, 0.0732 mmol), and (*R*)-MTPA-Cl (9.2 µL, 0.0488 mmol) were added, and stirring was continued at room temperature for an additional 26 h. The reaction was quenched with water, and the mixture was extracted with ethyl acetate. The organic phase was washed with brine and dried over anhydrous $MgSO_4$. The crude product was purified by silica gel column chromatography (Wako Gel® C-200, 50% ethyl acetate/*n*-hexane) to afford **S24** as a colorless oil (6.0 mg, quant.).

Based on the $^1$H-NMR spectra of **S23** and **S24**, the chemical-shift differences were calculated as $\Delta\delta = \delta((S)\text{-ester}) - \delta((R)\text{-ester})$. Negative $\Delta\delta$ values were observed for protons **a** and **b**, whereas positive $\Delta\delta$ values were obtained for protons **c**, **d**, and **e**. This pattern established that **MT1** possesses the intended *R* configuration (See Scheme S2).

***Outline of the synthesis of [$^{125}$I]pMT***

The synthetic route to **pMT** is outlined in Scheme **S3**. MOM protection of **S6** afforded **S25**, which was subsequently subjected to removal of the TBDPS protecting group and Parikh–Doering oxidation to give aldehyde **S27**. An Evans aldol reaction of **S27** with the chiral auxiliary **S17** afforded **S28**. Protection of the hydroxy group of **S28** as a TBS ether gave **S29**, which was subsequently reduced with DIBAL-H to afford **S30**. **S31** was synthesized from **S30** via a Horner–Wadsworth–Emmons reaction with $Ph_3P^+CH_2I\cdot I^-$. Following removal of the MOM protecting group, **S33** was synthesized by a Stille coupling reaction with **S16**. Mitsunobu reaction between **S33** and **S34** (prepared in *ref. S4*) afforded **S35**, and subsequent removal of the TBS protecting group gave **pMT**. **S36** was prepared from **S35** by tributylstannylation. Following removal of the TBS protecting group, the resulting stannyl precursor **S37** was converted to **[$^{125}$I]pMT** using $H_2O_2$ as the oxidizing agent and [$^{125}$I]NaI as the radioiodine source (*ref. S5*).

**Scheme S3.**

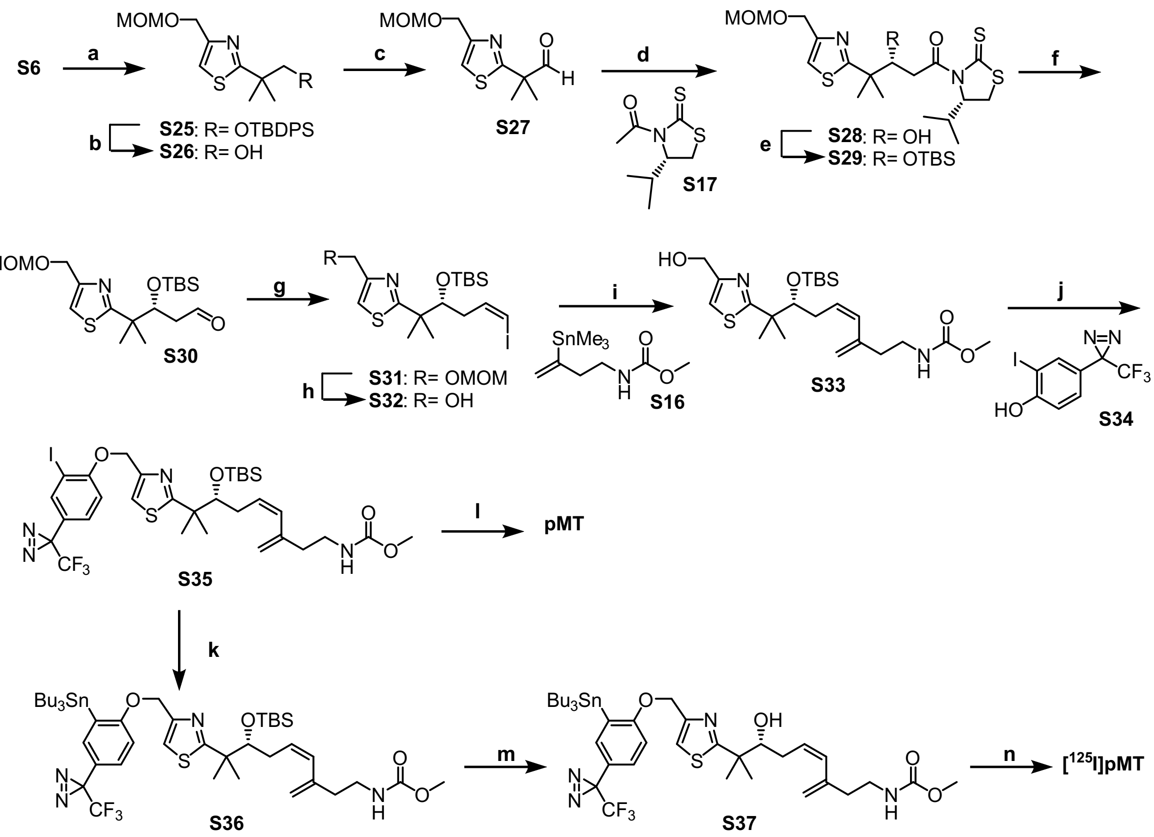


*Reagents and conditions*; (a) MOMCl, *i*$Pr_2NEt$, $CH_2Cl_2$, 0 ˚C to rt, 3 h, quant.; (b) TBAF·$3H_2O$, THF, 55°C, 22 h, 92%; (c) pyridine·$SO_3$, $Et_3N$, DMSO, $CH_2Cl_2$, 0 ˚C to rt, 2 h, 89%; (d) **S17**, $TiCl_4$, *i*$Pr_2NEt$, $CH_2Cl_2$, -50°C, 2 h, 75%; (e) TBSOTf, 2,6-lutidine, $CH_2Cl_2$, 0 ˚C to rt, 2 h, 71%; (f) DIBAL-H, toluene, -78 ˚C, 1 h, 84%; (g) $Ph_3P^+CH_2I{\cdot}I^-$, NaHMDS, HMPA, THF, -78°C, 20 min, 79%; (h) 10% AcCl/MeOH, rt, 24 h, 75%; (i) **S16**, CuI, $Pd(Ph_3P)_4$, CsF, DMF, rt, 40 min, 70%; (j) **S34**, DIAD, $PPh_3$, THF, rt, 2 h, 56%; (k) TBAF·$3H_2O$, THF, 45°C, 4 h, 36%; (l) $(SnBu_3)_2$, $Pd(CH_3CN)_2Cl_2$, HMPA, rt, 30 min; (m) TBAF·$3H_2O$, THF, 45°C, 2 h, 8% (2 steps); (n) [$^{125}$I]NaI, 0.02 M aqueous HCl, 30% (w/v) aqueous $H_2O_2$, 3 min, 26%.

**Synthesis of S26:** To an ice-cooled solution of **S6** (560 mg, 1.32 mmol) and *i*-$Pr_2NEt$ (898 μL, 5.28 mmol) in $CH_2Cl_2$ (45 mL) under a nitrogen atmosphere, MOM-Cl (1.00 mL, 13.2 mmol) was added. The mixture was allowed to warm to room temperature and stirred for 3 h. The reaction was quenched with saturated aqueous $NH_4Cl$, and the mixture was extracted with $CH_2Cl_2$. The organic phase was washed with water and brine, dried over anhydrous $MgSO_4$, and concentrated *in vacuo* to afford crude **S25**.

The MOM-protected alcohol **S25** was dissolved in THF (5 mL), and TBAF·$3H_2O$ (1.5 g, 4.95 mmol) was added. The mixture was stirred for 22 h at 55 °C, quenched with water, and extracted with ethyl acetate. The organic phase was washed with water and brine and dried over anhydrous $Na_2SO_4$. The crude product was purified by silica gel column chromatography (Wako Gel® C-200, 30–70% ethyl acetate/*n*-hexane) to afford **S26** as a colorless oil (280 mg, 1.24 mmol, 94% over two steps): $^1$H-NMR (400 MHz, $CDCl_3$) $\delta$ 7.12 (s, 1H), 4.73 (s, 2H), 4.66 (s, 2H), 3.70 (s, 2H), 3.40 (s, 3H), 1.40 (s, 6H); $^{13}$C-NMR (125 MHz, $CDCl_3$) $\delta$ 179.93, 152.84, 114.96, 96.23, 71.72, 65.21, 55.63, 42.43, 26.58 (2C).

**Synthesis of S27:** To an ice-cooled solution of **S26** (410 mg, 1.75 mmol) in $CH_2Cl_2$ (7 mL) under a nitrogen atmosphere, $Et_3N$ (974 μL, 7.00 mmol) and pyridine·$SO_3$ (1.10 g, 7.00 mmol, dissolved in 7 mL of DMSO) were added. The mixture was stirred at room temperature for 3 h and then quenched with saturated aqueous $NaHCO_3$. After removal of $CH_2Cl_2$ *in vacuo*, the resulting residue was extracted with $Et_2O$. The organic phase was washed with saturated aqueous $NH_4Cl$ and brine and dried over anhydrous $MgSO_4$. The crude product was purified by silica gel column chromatography (Wako Gel® C-200, 40% ethyl acetate/*n*-hexane) to afford **S27** as a colorless oil (360 mg, 1.55 mmol, 89%): $^1$H-NMR (400 MHz, $CDCl_3$) $\delta$ 9.68 (s, 1H), 7.21 (s, 1H), 4.73 (S, 2H), 4.69 (s, 2H), 3.70 (s, 2H), 3.40 (s, 3H), 1.57 (s, 6H); $^{13}$C-NMR (100 MHz, $CDCl_3$) $\delta$ 200.09, 172.51, 154.32, 116.20, 96.18, 65.19, 55.62, 52.03, 23.14 (2C); ESI-MS *(m/z)* 230.1 $[M+H]^+$.

**Synthesis of S28:** To a solution of **S17** (470 mg, 2.32 mmol) in $CH_2Cl_2$ (8 mL), $TiCl_4$ (270 μL, 2.48 mmol) was added at −50 °C under a nitrogen atmosphere, and the mixture was stirred at -50 °C for 20 min. *i*-$Pr_2NEt$ (422 μL, 2.48 mmol) was then added, and stirring was continued at -45 °C for 1 h. The reaction mixture was cooled to -78 °C, and a solution of **S27** (360 mg, 1.55 mmol) in $CH_2Cl_2$ (5 mL) was added. The mixture was stirred at -40 °C for 3 h and then quenched with phosphate buffer and Rochelle salt solution. The mixture was extracted with $CH_2Cl_2$, and the organic phase was washed with brine and dried over anhydrous $MgSO_4$. The crude product was purified by silica gel column chromatography (Wako Gel® C-200, 20–70% ethyl acetate/*n*-hexane) to afford **S28** as a yellow oil (500 mg, 1.16 mmol, 75%): $^1$H-NMR (400 MHz, $CDCl_3$) $\delta$ 7.26 (s, 1H), 5.12 (dd, $J$ = 6.8, 6.8 Hz, 1H), 4.72 (s, 2H), 4.67 (s, 2H), 4.34 (brt, 1H),

3.56-3.45 (m, 2H), 3.40 (d, $J$ = 18.7 Hz, 2H), 3.40 (s, 3H), 3.00 (d, $J$ = 11.4 Hz, 1H), 2.45-2.30 (m, 1H), 1.46 (s, 3H), 1.44 (s, 3H), 1.04 (d, $J$ = 6.8 Hz, 3H), 0.96 (d, $J$ = 7.0 Hz, 3H); $^{13}$C-NMR (100 MHz, $CDCl_3$) $\delta$ 203.39, 178.72, 172.80, 153.35, 115.02, 96.17, 75.12, 72.01, 65.26, 55.62, 44.63, 40.98, 31.03, 30.80, 26.61, 24.78, 19.28, 17.90, 14.38; ESI-MS (*m/z*) 433.2 $[M+H]^+$.

**Synthesis of S29:** To an ice-cooled solution of **S28** (500 mg, 1.16 mmol) in $CH_2Cl_2$ (10 mL), 2,6-lutidine (230 µL, 1.97 mmol) and TBSOTf (400 µL, 1.74 mmol) were added under a nitrogen atmosphere. The mixture was stirred at 0 °C for 30 min and then quenched with saturated aqueous $NH_4Cl$. The mixture was extracted with $Et_2O$, and the organic phase was washed successively with Rochelle salt solution, water, and brine and dried over anhydrous $MgSO_4$. The crude product was purified by silica gel column chromatography (Wako Gel® C-200, 15% ethyl acetate/*n*-hexane) to afford **S29** as a yellow oil (430 mg, 0.794 mmol, 71%): $^1$H-NMR (400 MHz, $CDCl_3$) $\delta$ 7.26 (s, 1H), 5.04 (t, $J$ = 7.0 Hz, 1H), 4.69 (t, $J$ = 4.9 Hz, 1H), 4.69 (s, 2H), 4.65 (s, 2H), 3.42-3.39 (m, 1H), 3.37 (s, 3H), 3.30 (d, $J$= 4.1 Hz, 2H), 2.97 (d, $J$ = 11.5 Hz, 1H), 2.40-2.20 (m, 1H), 1.41 (s, 3H), 1.34 (s, 3H), 1.00 (d, $J$ = 6.8 Hz, 3H), 0.91 (d, $J$ = 7.0 Hz, 3H), -0.07 (s, 6H); $^{13}$C-NMR (100 MHz, $CDCl_3$) $\delta$ 202.72, 177.80, 172.08, 152.82, 115.11, 95.95, 75.33, 71.60, 65.30, 55.62, 44.46, 43.48, 30.98, 30.80, 26.06 (3C), 25.78, 25.49, 19.19, 18.34, 18.08, -4.41, -5.00; ESI-MS (*m/z*) 547.3 $[M+H]^+$.

**Synthesis of S31:** To a solution of **S29** (150 mg, 0.274 mmol) in toluene (3 mL), DIBAL-H (in *n*-hexane, 686 µL, 0.686 mmol) was added dropwise at -78 °C under an argon atmosphere, and the mixture was stirred at -78 °C for 1 h. The reaction was quenched with aqueous Rochelle salt solution at -78 °C and allowed to warm to room temperature, followed by stirring for 1 h. The mixture was extracted with $Et_2O$, and the organic phase was washed with Rochelle salt solution and brine and dried over anhydrous $MgSO_4$. The crude product was purified by silica gel column chromatography (Wako Gel® C-200, 20% ethyl acetate/*n*-hexane) to afford **S30** (86 mg, crude), which was used directly in the next step without further purification.

To a solution of $Ph_3P^+CH_2I{\cdot}I^-$ (430 mg, 0.81 mmol) in THF (2 mL), NaHMDS (810 µL, 1.62 mmol) was added under an argon atmosphere, and the mixture was stirred at room temperature for 5 min. The reaction mixture was cooled to -78 °C, and HMPA (738 µL, 4.24 mmol) was added. Stirring was continued at -78 °C for an additional 5 min, followed by the addition of **S30** (86 mg, crude). The mixture was stirred at -78 °C for 20 min and then quenched with water. The reaction mixture was allowed to warm to room temperature and extracted with $Et_2O$. The organic phase was washed with water and brine and dried over anhydrous $MgSO_4$. The crude product was purified by silica gel column chromatography (Wako Gel® C-

200, 5–20% ethyl acetate/*n*-hexane) to afford **S31** as a pale yellow oil (90 mg, 0.176 mmol, 66% over two steps): $^{1}$H-NMR (400 MHz, $CDCl_3$) $\delta$ 7.09 (s, 1H), 6.20-6.05 (m, 2H), 4.74 (s, 2H), 4.69 (s, 2H), 4.17 (t, $J$ = 5.3 Hz, 1H), 3.41 (s, 3H), 1.44 (s, 3H), 1.39 (s, 3H), 0.87 (s, 9H), 0.04 (s, 3H), -0.11 (s, 3H); $^{13}$C-NMR (100 MHz, $CDCl_3$) $\delta$ 178.68, 152.99, 138.85, 115.10, 96.04, 83.01, 78.03, 65.41, 55.63, 46.45, 39.58, 26.19 (3C), 25.90, 24.97, 18.38, -3.58, -4.54; ESI-MS (*m/z*) 512.2 $[M+H]^+$.

**Synthesis of S32:** To an ice-cooled solution of **S31** (30 mg, 0.059 mmol) in $CH_2Cl_2$ (2 mL), 10% AcCl in MeOH (1 mL) was added, and the mixture was stirred at room temperature for 26 h. The reaction was quenched with saturated aqueous $NaHCO_3$, and the mixture was extracted with ethyl acetate. The organic phase was washed with brine and dried over anhydrous $MgSO_4$. The crude product was purified by silica gel column chromatography (Wako Gel® C-200, 20–40% ethyl acetate/*n*-hexane) to afford **S32** as a colorless oil (21 mg, 0.045 mmol, 77%): $^{1}$H-NMR (400 MHz, $CDCl_3$) $\delta$ 7.03 (s, 1H), 6.15-6.05 (m, 2H), 4.73 (s, 2H), 4.69 (s, 2H), 4.14 (t, $J$ = 5.2 Hz, 1H), 2.35-2.20 (m, 2H), 1.44 (s, 3H), 1.39 (s, 3H), 0.87 (s, 9H), 0.04 (s, 3H), -0.08 (s, 3H); $^{13}$C-NMR (100 MHz, $CDCl_3$) $\delta$ 178.94, 155.52, 138.68, 113.92, 83.11, 78.05, 61.33, 46.47, 39.55, 26.18 (3C), 26.02, 25.02, 18.39, -3.57, -4.51, 178.94; ESI-MS (*m/z*) 468.2 $[M+H]^+$.

**Synthesis of S33:** A solution of **S32** (40 mg, 0.092 mmol) and **S16** (48 mg, 0.165 mmol) in DMF (0.5 mL) was purged with argon. CuI (3 mg, 0.016 mmol), $Pd(PPh_3)_4$ (6 mg, 0.005 mmol), and CsF (14 mg, 0.092 mmol) were then added successively, and the mixture was stirred at room temperature for 10 min. The reaction was quenched with water, and the mixture was extracted with $Et_2O$. The organic phase was washed with water and dried over anhydrous $MgSO_4$. The crude product was purified by silica gel column chromatography (Wako Gel® C-200, 20–50% ethyl acetate/*n*-hexane) to afford **S33** as a yellow oil (30 mg, 0.064 mmol, 70%): $^{1}$H-NMR (400 MHz, $CDCl_3$) $\delta$ 7.04 (s, 1H), 5.87 (d, $J$ = 9.2 Hz, 1H), 5.60~5.70 (m, 1H), 4.95 (s, 1H), 4.90 (brs, 1H), 4.82 (s, 1H), 4.72 (s, 2H), 3.99 (t, $J$ = 5.3 Hz, 1H), 3.65 (s, 3H), 3.35-3.25 (m, 2H), 2.46 (t, $J$ = 6.6 Hz, 2H), 2.21 (t, $J$ = 6.7 Hz, 2H), 1.41 (s, 3H), 1.36 (s, 3H), 0.87 (s, 9H), 0.04 (s, 3H), -0.05 (s, 3H); $^{13}$C-NMR (100 MHz, $CDCl_3$) $\delta$ 178.94, 157.31, 155.69, 142.20, 130.61, 129.41, 123.89, 116.02, 113.84, 79.63, 52.21, 46.48, 39.80, 37.38, 26.42, 26.16, 24.83, 18.43, -3.38, -4.37; ESI-MS (*m/z*) 469.3 $[M+H]^+$.

**Synthesis of S34:** This compound was synthesized according to the procedure described in *ref. S4*.

**Synthesis of S35:** A solution of **S33** (14 mg, 0.03 mmol), **S34** (30 mg, 0.06 mmol), and $PPh_3$ (32 mg, 0.06 mmol) in THF (0.1 mL) was stirred at room temperature for 15 min. The mixture was then cooled to 0 °C, and DIAD (63 μL, 0.06 mmol) was added. Stirring was continued at room temperature for 1 h. The reaction mixture was concentrated *in vacuo*, and the residue was purified by silica gel column chromatography (Wako Gel® C-200, 0–50% ethyl acetate/*n*-hexane) to afford **S35** as a colorless oil (13 mg, 0.017 mmol, 56%): $^1$H-NMR (400 MHz, $CDCl_3$) $\delta$ 7.60 (d, $J$ = 2.0 Hz, 1H), 7.37 (s, 1H), 7.20 (dd, $J$ = 2.0, 6.4 Hz, 1H), 6.91 (dd, $J$ = 3.4, 5.2 Hz, 1H), 5.66 (d, $J$ = 11.6 Hz, 1H), 5.55-5.45 (m, 1H), 5.24 (s, 1H), 4.97 (s, 1H), 4.84 (s, 1H), 4.72 (s, 2H), 4.01 (t, $J$ = 5.0 Hz, 1H), 3.64 (s, 3H), 3.21 (m, 2H), 2.46 (t, $J$ = 6.6 Hz, 2H), 2.45-2.25 (m, 2H), 2.23 (t, $J$ = 6.3 Hz, 2H), 1.42 (s, 3H), 1.37 (s, 3H), 0.88 (s, 9H), 0.05 (s, 3H), -0.05 (s, 3H); $^{13}$C-NMR (100 MHz, $CDCl_3$) $\delta$ 179.25, 158.49, 157.21, 156.81, 150.51, 142.11, 137.98, 130.70, 129.40, 123.44, 122.53 (q, $J_{CF}$ = 273 Hz), 116.09, 115.85, 112.41, 86.14, 79.62, 68.06, 52.27, 46.67, 39.61, 37.58, 33.42, 28.95 (q, $J_{CF}$ = 38 Hz), 27.50 (3C), 26.11, 25.14, -3.36, -4.35; ESI-MS (*m/z*) 779.2 $[M+H]^+$.

**Synthesis of pMT:** To an ice-cooled solution of **S35** (13 mg, 0.0167 mmol) in THF (1.0 mL), $TBAF\cdot 3H_2O$ (26 mg, 0.0835 mmol) was added, and the mixture was stirred at 45 °C for 4 h. The reaction was quenched with water, and the mixture was extracted with ethyl acetate. The organic phase was washed with brine and dried over anhydrous $Na_2SO_4$. The crude product was purified by silica gel column chromatography (Wako Gel® C-200, 10–20% ethyl acetate/*n*-hexane) to afford **pMT** as a colorless oil (4 mg, 0.006 mmol, 36%): $^1$H-NMR (400 MHz, $CDCl_3$) $\delta$ 7.60 (d, $J$ = 2.0 Hz, 1H), 7.37 (s, 1H), 7.20 (dd, $J$ = 2.0, 6.4 Hz, 1H), 6.91 (dd, $J$ = 3.4, 5.2 Hz, 1H), 5.88 (d, $J$ = 11.4 Hz, 1H), 5.70-5.60 (m, 1H), 5.23 (s, 1H), 5.01 (s, 1H), 4.86 (s, 1H), 4.25 (brs, 1H), 3.80 (m, 1H), 3.64 (s, 3H), 3.35-3.10 (m, 2H), 2.50-2.20 (m, 4H), 1.46 (s, 3H), 1.43 (s, 3H); $^{13}$C-NMR (100 MHz, $CDCl_3$) $\delta$ 180.53, 158.34, 157.34, 150.80, 142.59, 138.01, 130.53, 128.64, 123.64, 122.53 (q, $J_{CF}$ = 273 Hz), 116.16, 115.59, 115.31, 112.38, 78.45, 77.43, 67.82, 52.13, 45.16, 39.62, 37.43, 30.90, 28.95 (q, $J_{CF}$ = 38 Hz), 26.65, 24.29; ESI-MS (*m/z*) 665.2 $[M+H]^+$.

**Synthesis of S36:** To a solution of **S35** (25 mg, 0.032 mmol) in HMPA (0.1 mL), $Sn_2Bu_6$ (48 μL, 0.0936 mmol) and $Pd(MeCN)_2Cl_2$ (3 mg, 0.013 mmol) were added under an argon atmosphere, and the mixture was stirred at room temperature for 1 h. The reaction mixture was directly applied to a silica gel column (Wako Gel® C-200) and chromatographed using 0–10% ethyl acetate/*n*-hexane as the eluent to afford **S36** (8 mg, containing inseparable $Sn_2Bu_6$-derived byproducts), which was used directly in the next step without further purification.

**Synthesis of S37:** To an ice-cooled solution of **S36** (8 mg, crude) in THF (0.5 mL), TBAF·$3H_2O$ (6.7 mg, 0.021 mmol) was added, and the mixture was stirred at 45 °C for 2 h. The reaction was quenched with water, and the mixture was extracted with ethyl acetate. The organic phase was washed with brine and dried over anhydrous $Na_2SO_4$. The crude product was purified by silica gel column chromatography (Wako Gel® C-200, 15–30% ethyl acetate/*n*-hexane) to afford **S37** as a colorless oil (2.2 mg, 0.0027 mmol, 8% over two steps): $^1$H-NMR (400 MHz, $CDCl_3$) $\delta$ 7.19 (s, 1H), 7.13 (d, $J$ = 2 Hz, 1H), 7.10 (d, $J$ = 6.4 Hz, 1H), 6.84 (dd, $J$ = 3.4, 5.2 Hz, 1H), 5.88 (d, $J$ = 12.2 Hz, 1H), 5.70-5.60 (m, 1H), 5.34 (brs, 1H), 5.13 (s, 1H), 5.02 (s, 1H), 4.87 (s, 1H), 4.28 (m, 1H), 3.83 (d, $J$ = 9.8 Hz, 1H), 3.62 (s, 3H), 3.35-3.10 (m, 2H), 2.50-2.10 (m, 4H), 1.55-1.35 (m, 12H), 1.35-1.20 (m, 6H), 1.10-0.90 (m, 6H); $^{13}$C-NMR (100 MHz, $CDCl_3$) $\delta$ 180.44, 163.72, 157.31, 151.49, 142.64, 135.38, 131.30, 130.60, 128.75, 122.53 (q, $J_{CF}$ = 273 Hz), 121.80, 116.10, 115.43, 110.04, 78.29, 77.42, 66.25, 52.08, 45.18, 39.57, 37.40, 30.79, 29.24, 28.95 (q, $J_{CF}$ = 38 Hz), 27.46, 26.65, 24.16, 13.87, 10.09.

**Synthesis of [$^{125}$I]pMT:** To a solution of **S37** (1.0 mM) in ethanol (20 µL) in a screw-capped 1.5-mL plastic tube, [$^{125}$I]sodium iodide (PerkinElmer, NEZ033A, 1 mCi, 2000 Ci/mmol, 10 µL) was added, and the mixture was vortexed. Radioiodination was initiated by the addition of 30% (w/v) aqueous $H_2O_2$, followed by vortexing and incubation at room temperature for 3 min. The reaction was terminated by the addition of 0.1 M aqueous NaOH (20 µL), and the resulting mixture was carefully extracted with $CHCl_3$ (100 µL × 3). The combined organic layers were concentrated using a vacuum centrifugal evaporator. The crude product was dissolved in MeOH (50 µL) and subjected to HPLC purification (Shimadzu LC-10AS, Kyoto, Japan) using an ODS column (COSMOSIL 5C18-MS-II, 4.6 × 150 mm, Nacalai Tesque, Kyoto, Japan) at a flow rate of 800 µL/min with an isocratic 87% MeOH/water system as the eluent. Fractions were collected every 30 s (400 µL each). To determine the elution profile of the radioactivity and assess radiochemical purity, each fraction was analyzed by γ-counting and radio-TLC. Fractions exhibiting strong radioactivity at the retention time corresponding to **pMT** were combined, and the solvent was removed using a vacuum centrifugal evaporator. **[$^{125}$I]pMT** was stored as an ethanolic solution (2.0 µM) at 4 °C. The radiochemical yield of **[$^{125}$I]pMT**, based on the initial amount of [$^{125}$I]NaI, was 26%.

**References (Synthesis only)**

S1. Sugiyama, H. Yokokawa, F. and Shioiri, T (2000) Asymmetric total synthesis of (-)-mycothiazole, *Org. Lett. 2*, 2149-2152.

S2. McNulty, J. and McLeod, D. (2017) Total enantioselective synthesis of the endophytic fungal polyketide phomolide H and its structural revision, *Eur. J. Org. Chem.* 2017, 29-33.

S3. McNulty, J. and McLeod, D. (2017) Total enantioselective synthesis of the endophytic fungal polyketide phomolide H and its structural revision, *Eur. J. Org. Chem.* 2017, 29-33.

S4. Uno, S., Masuya, T., Zdorevskyi, O., Ikunishi, R., Shinzawa-Itoh, K., Lasham, J., Sharma, V., Murai, M., and Miyoshi, H. (2022) Diverse reaction behaviors of artificial ubiquinones in mitochondrial respiratory complex I. *J. Biol. Chem.* 298, 102075.

S5. Nishida, M., Pecorilla, C., Masuya, T., Hirano, K., Abe, M, Zdorevskyi, O., Sharma V., Miyoshi, H., and Murai, M. (2025) Dynamic binding of acetogenin-type inhibitors to mitochondrial complex I revealed by photoaffinity labeling, *Biochim. Biophys. Acta (Bioenergetics)*, 1866, 149566.
